# Dissecting glial scar formation by spatial point pattern and topological data analysis

**DOI:** 10.1101/2023.10.04.560910

**Authors:** Daniel Manrique-Castano, Dhananjay Bhaskar, Ayman ElAli

## Abstract

Glial scar formation represents a fundamental response to central nervous system (CNS) injury. It is mainly characterized by a well-defined spatial rearrangement of reactive astrocytes and microglia. The mechanisms underlying glial scar formation have been extensively studied, yet quantitative descriptors of the spatial arrangement of reactive glial cells remain limited. Here, we present a novel approach using point pattern analysis (PPA) and topological data analysis (TDA) to quantify spatial patterns of reactive glial cells after experimental ischemic stroke in mice. We provide open and reproducible tools using *R* and *Julia* to quantify spatial intensity, cell covariance and conditional distribution, cell-to-cell interactions, and short/long-scale arrangement, which collectively disentangle the arrangement patterns of the glial scar. This approach unravels a substantial divergence in the distribution of reactive astrocytes and microglia after injury that conventional analysis methods cannot fully characterize. PPA and TDA are valuable tools for studying the complex spatial arrangement of reactive glia and other nervous cells following CNS injuries and have potential applications for evaluating glial-targeted restorative therapies.

## Introduction

Glial scar formation represents a crucial response to central nervous system (CNS) injuries, such as traumatic brain injury and ischemic stroke (Bradbury and Burnside 2019). The scar is primarily composed of reactive astrocytes, activated microglia, and polydendrocytes/NG2^+^ cells, which rearrange spatially to establish a demarcated, yet dynamic, barrier around the lesioned tissue (Wanner et al. 2013; Hackett and Lee 2016; Bellver-Landete et al. 2019; Zhang et al. 2022). This enables compartmentalization of the injured tissue (Conforti et al. 2022) to limit the spread of toxic and inflammatory mediators towards healthy regions (Voskuhl et al. 2009; Yoshizaki et al. 2021). The organization of reactive glia affects the structural and functional integrity of the lesioned tissue, making processes such as axonal sprouting and neurovascular repair a vital target for therapeutic interventions (Anderson et al. 2016; Fu et al. 2020).

In recent years, *in vivo* and *in vitro* models of CNS injuries have identified several molecular and cellular markers associated with scar formation (Yang et al. 2020), including glial fibrillary acid protein (GFAP) and ionized calcium-binding adaptor molecule-1 (IBA1) (Kamphuis et al. 2015; Schacke et al. 2022; Lu et al. 2021). Together with the massive arrangement of reactive glial cells in the infarct core, these markers presage scar formation in various CNS injuries, including ischemic stroke (Buscemi et al. 2019; Manrique-Castano et al. 2021; Ito et al. 2001). However, current methods for analyzing glial scar formation primarily rely on qualitative assessments that lack spatiotemporal resolution and limit the examination of cellular arrangement patterns.

To address these limitations, we developed a comprehensive implementation of point pattern analysis (PPA) (Parra 2021; Jafari-Mamaghani, Andersson, and Krieger 2010; Davis et al. 2017), spatial clustering (Prodanov, Nagelkerke, and Marani 2007), and topological data analysis (TDA) (Bhaskar, Zhang, and Wong 2021; Masoomy et al. 2021; Bonilla, Carpio, and Trenado 2020) for the assessment of the spatiotemporal dynamics of glial scar formation after experimental ischemic stroke. PPA allows to determine the distribution patterns of objects in a given observation window through different metrics. It has multiple applications in ecology (Malavasi et al. 2023), epidemiology (Scarpone et al. 2020), and cancer biology (Maisel et al. 2022; Kaufmann et al. 2021). Conversely, TDA uses the tools of algebraic topology to extract patterns from complex data sets or point clouds, which may be difficult to see using traditional methods. This includes shape, structure, and connectivity (Vipond et al. 2021; Lawson et al. 2019). A central method in TDA is *persistent homology* (PH), which tracks the evolution of topological features, such as connected components, loops, voids, and higher-dimensional holes (Amézquita et al. 2020; Townsend et al. 2020). Recently, TDA has been suitable for modeling collective cell motion and migration (Bhaskar, Zhang, and Wong 2021; Bonilla, Carpio, and Trenado 2020), neuronal morphology (Kanari et al. 2017), and branching patterns in vascularization (Nardini et al. 2021). However, no direct applications of PPA/TDA exist for analyzing the dynamics of cell rearrangement following brain injury, which is critical for assessing scar formation and tissue integrity.

Our approach provides reproducible pipelines to quantify cell covariance, cell-to-cell interactions, and short and long-scale cumulative cell distributions. This allowed us to identify the divergent allocation of reactive astrocytes and activated microglia/macrophages, revealing a double-layered glial scar with unique dynamics throughout the injury that eludes the scrutiny of traditional methods. Furthermore, we build a machine-learning model to predict the time post-ischemia based on cell positions. We provide the raw data and annotated scripts for R and Julia in a GitHub repository to ensure replicability and encourage using PPA and TDA-based analysis for glial reactivity in other disease models.

## Materials and Methods

We share the raw microscopy images, data tables (10.5281/zenodo.8399976), and annotated scripts for the full reproducibility of our PPA-TDA analysis (https://github.com/elalilab/GlialScar_PPA-TDA_2022). The companion Quarto notebook (QN) in the Github repository details all data handling, statistical models, and analysis pipelines. See also the Open Science Framework repository (OSFr) for additional material (DOI 10.17605/OSF.IO/3VG8J).

### Animals and experimental approach

The animals were housed in groups of 3-5 per cage and acclimated to standard laboratory conditions (12 hours light/dark cycle; Lights on at 7:00 AM and off at 7:00 PM) with free access to chow and water. Six-month-old C57BL/6J mice (a total of 38) were subjected to focal transient cerebral ischemia via middle cerebral artery occlusion (MCAo), with an additional group of age/weight-matched mice (n = 5) included as a control/reference. Ten animals either died after MCAo or were euthanized according to veterinary recommendations, and two animals that did not exhibit signs of cerebral ischemia were excluded from the analysis. Animal survival at 5, 15, and 30 days post-ischemia (DPI) was randomized using the Research Randomizer online tool (https://www.randomizer.org/). The experimental procedure is summarized in Suppl. Figure 1A. All animal procedures and handling were performed according to the Canadian Council on Animal Care guidelines, as implemented by *Comité de Protection des Animaux de l’Université Laval-3* (CPAUL-3; Protocol # 20-470).

### Middle cerebral artery occlusion (MCAo)

Transient focal ischemic stroke was induced by 30 min MCAo to produce a cortical-striatal lesion in the territory of the MCA (see Suppl. methods). Mice were anesthetized with 1.5% v/v isoflurane and the rectal temperature was maintained at 37.0°C using a feedback-controlled heating system. A longitudinal midline incision was made to expose the left common carotid artery (CCA). We permanently ligated the lowest visible segment of the CCA and placed a temporal loose ligature anterior to the bifurcation giving rise to the external carotid artery (ECA) and the internal carotid artery (ICA). Ligation of the ECA was avoided to prevent masticatory lesions and to improve animal welfare after ischemia (Dittmar et al. 2005). An incision was then made between the permanent and temporal ligatures to insert a silicone resin-coated nylon monofilament (MCAO suture, Doccol Corporation, Cat# 7022910PK5Re). The monofilament was advanced into the CCA to block the MCA fork at the polygon of Willis and tied tightly to prevent retraction. After 30 minutes, the monofilament was removed through the entry incision and the upper segment of the CCA was permanently ligated. Finally, the cervical incision was sutured and anesthesia was discontinued. Postoperative care was provided in the home cage by subcutaneous injection of 1 mL Ringer’s lactate twice daily for at least three days or until the preoperative weight was regained. We provided wet food and diet gel boost (DietGel® Boost, Clear H2O, Cat #72-04-5022) in the bottom of the cage to facilitate feeding.

### Brain sectioning and immunolabeling

Mice were euthanized by transcardial perfusion with 0.9% sodium chloride (NaCl) followed by 4% paraformaldehyde (PFA). The brains were removed and post-fixed by submersion in 4% PFA during 16-24h. After rinsing and cryoprotection, brains were frozen and stored at −80°C until sectioning with a freezing microtome (Leica Biosystems, ON, Canada) (see Suppl. methods). Thirty μm thick coronal sections were cut and mounted serially (Suppl. Table 1) onto SuperFrost^®^ Plus slides (Fisher Scientific, ON, Canada). We performed routine immunofluorescence staining with antibodies against IBA1, NeuN, GFAP, and DAPI (see Suppl. methods).

### Brain imaging

We scanned 6-7 whole brain coronal sections per animal (Suppl. Table 1) at 5x magnification using an AxioScan Z1 slide scanner (Carl Zeiss Canada, ON, Canada), with the parameters specified in Suppl. Table 2. To enhance cell detection, we processed the multichannel images in FIJI (Schindelin et al. 2012) using the scripts available on our GitHub repository. We manually rotated or flipped the images (when necessary) and removed interfering brain pieces/artifacts to facilitate alignment with the Allen reference atlases. Please refer to Suppl. Figure 1B for a detailed view.

To examine neuronal and glial distribution at the level of the MCA territory, we imaged section 3 (bregma, 0.44 to −0.06) at 10x magnification using a Zeiss Axio Observer.Z1 inverted epifluorescence microscope. The acquisition parameters are listed in Suppl. Table 3. We acquired a horizontal ROI from the ventricular area to the outer border of the dorsolateral cerebral cortex, as depicted in Suppl. Figure 1C. To delineate the observation windows for PPA, we sharpened the sections using FIJI and manually removed blank regions.

### Slice alignment, cell detection, and quantification

We aligned the 5x magnification whole brain slices to the Allen Mouse Brain Atlas using the Aligning Big Brains & Atlases (ABBA) plugin for FIJI, which performs automated 2D affine and spline in-plane registrations of brain slices. We used the BigWarp-framed tool to correct slice orientation and registration when required. The aligned/registered brain slices were then exported to QuPath (Bankhead et al. 2017) to perform unbiased cell detection and quantification of NeuN-, GFAP-, and IBA1-expressing cells in the cortex (CTX), nuclei (CNU), fiber tracts, midbrain (MB), and inter-brain (IB) using QuPath scripts (see GitHub repository). In particular, we used a watershed algorithm and object classifiers to select individual nuclei with normal NeuN immunolabeling. In contrast, GFAP^+^ and IBA1^+^ cells were thresholded for marker expression typical of reactive cells. Due to irregular glial morphology and variable size, unbiased detection may include whole or stained cell fragments. We also applied comparable cell detection and quantification to 10x magnification images at the level of the MCA territory, without alignment to the Allen brain atlas.

In this article, we refer to GFAP^+^ cells as reactive astrocytes, recognizing that progenitors can also express this marker in cell niches such as the ventricular zone. Similarly, we refer to IBA1^+^ cells as activated microglia, knowing that this protein is also expressed by some infiltrating immune cells in intra-lesional areas. This aspect does nor impact our conclusions since infiltraed IBA1^+^ cells fulfill similar functions as resident IBA1^+^ cells.

### Point pattern analysis (PPA)

We handled cell counts and coordinates obtained from QuPath using R/Rstudio. The supplementary QN fully describes the PPA approach using functions from the *spatstat* R-package (Baddeley, Rubak, and Turner 2015) with step-by-step implementation. The analyzed point patterns are available in the Zenodo repository. We performed PPA on section # 3 (bregma 0.44 to −0.06) of each brain, which includes the MCA territory imaged at 10x. We downsampled to 10% of the detected cells to improve computational efficiency and visualization.

Our PPA approach entails (1) the calculation of the *spatial intensity* (QN, section 1) to identify areas with divergent cell density and the assessment of the distribution of each cell type across the glial scar. (2) *Quadrant counts* to investigate the spatial relationship between different cell types using tessellations— a partition of space into non-overlapping regions (QN, section 5-6). (3) *rhohat* and multiple point process models (mppm) to calculate the relative/conditional intensity (see corresponding QN chapters). These nonparametric mppm allow us to estimate spatial distributions without making underlying assumptions. (4) Rasterization of the point patterns to create matrices of intensity-based gridded data to perform dispersion and distance measurements (Hijmans 2023) (QN, Section 7). (5) Calculation of the inhomogeneous *L-function*, which is Besag’s transformation of Ripley’s K-function, to assess cell-to-cell interactions within point patterns and deviations from complete spatial randomness (CSR) (QN, Section 9).

### Topological data analysis (TDA)

To calculate the PH to point cloud data, we compute Vietoris-Rips filtration from cell positions using the GPU-enabled *Ripser++* package (Zhang, Xiao, and Wang 2020). Additionally, we used a topological machine learning based approach (Hensel, Moor, and Rieck 2021) to predict DPI based on cell positions. We first applied persistent homology analysis to the point clouds to obtain topological features of the data.

Subsequently, we used a supervised learning algorithm, support vector regression (SVR) (Ben-Hur et al. 2008) with a radial basis function (RBF) kernel, to predict DPI. We evaluated the performance of our approach using a 10-fold cross-validation procedure (Dutschmann et al. 2023; Jung 2017). In each fold, we randomly split the dataset into training and testing sets, with 80% of the data used for training and 20% for testing. We repeated this procedure 10 times, with a different random split each time, and computed the mean and standard deviation of the prediction error across the 10 folds.

### Statistical analysis

In this study, we performed Bayesian modeling and statistical inference using the R-package *brms* (Bürkner 2018) and non-parametric modeling with *spatstat (Baddeley and Turner 2005; Baddeley, Rubak, and Turner 2015)*. The parameters are tailored to each dataset and specifically defined in the QN, together with model validation. The results of each model are reported in Suppl. Tables 4-11.

We used R-packages *ggplot* and ggdist to visualize estimates and their uncertainty. Density kernels, tessellations, *rhohat*, and L-functions were plotted using *spatstat*-linked R-base plotting system. We present the complete posterior distributions with half-eye densities and point intervals. The contrast between time points (half-eye and point intervals) is evaluated based on a region of practical equivalence (ROPE), which indicates the region falling within the intragroup variance estimated by the fitted models. We estimated the contrast between time points using the *emmeans* package and used the *hypothesis* function from *brms* to obtain point estimates and 95% credible intervals (CI). Note that Bayesian CIs differ from confidence intervals (Hespanhol et al. 2019). We performed scientific inference using the entire posterior distribution and calculated the probability of falling within the ROPE using the entire posterior.

For conditional distribution estimates (rhohat), we plot the pooled (per group) mean (magenta) with two-sigma confidence intervals (shaded gray region). In addition, we plotted the pooled mean intensity (dashed cyan), and the lower (solid black) and upper (dotted green) limits of the two-sigma confidence intervals for the mean intensity. The scales of the plots were weighted or constrained for optimal visualization, depending on the available ranges for the computed function. For images of the MCA territory at 10x, we plotted L-functions with 95% confidence intervals (shaded gray area) and performed a studentized permutation test (Hahn 2012) using the *studpermu.test* function from *spatstat*.

For TDA, we used Betti curves to represent the Betti numbers of the dataset against a parameter, such as a threshold or a scale. We analyzed the Betti curves using the non-parametric two-sample Kolmogorov-Smirnov (KS) test to compare the cumulative distribution functions (CDFs). We also employed permutation tests to randomly permute the labels of the data points and compute the Betti curves for each permutation. We repeated this process many times to obtain a null distribution of the test statistic.

## Results

We performed PPA and TDA on NeuN^+^ (neurons), GFAP^+^ (reactive astrocytes), and IBA1^+^ (activated microglia) to quantify the spatiotemporal and topological arrangement of glial scars after ischemic stroke. Our approach provides essential quantitative insights into the spatial intensity (the cell density fluctuations) and the relative distribution of glial cells (mapped by point patterns) with respect to different covariates. Furthermore, we demonstrate that TDA enables researchers to predict stages of injury and the distribution of different cell groups based on the allocation of single cell types. Raw data, data handling, statistical models, and model validation are available on-line for full reproducibility of our results.

### Spatial intensity of neurons and glial cells following injury

Ischemic stroke is associated with profound changes in the density of neurons and glial cells at the lesioned tissue over time (Sofroniew and Vinters 2009). Therefore, we employed PPA to examine the variation of the spatial intensity of NeuN^+^, GFAP^+^, and IBA1^+^ cells during the course of injury. Details of the raw data, model selection, and diagnostics are available (QN, Section 2 and Suppl. Table 4).

The spatial intensity of NeuN^+^ cells showed a marked reduction of 17 units (95% CI = 9.7 - 24.7) at 5 DPI (Figure 1C, top). No substantial changes were observed from this stage to 15 DPI (−1.64, 95% CI = −8.6 - 5.4, ROPE = 0.89). However, our results show that at 30 DPI, the spatial intensity of NeuN^+^ cells exhibits a partial recovery of 7 units (95% CI = 1.24 - 13.7, ROPE = 0.54). This output may be driven by increased neuronal density resulting from severe brain shrinkage or the partial recovery of NeuN immunoreactivity associated with the restauration of metabolism in the penumbra (Unal-Cevik et al. 2004). Thus, the analysis of spatial intensity captures the complex variations in neuronal density associated with brain shrinkage and cell antigenicity.

**Figure 1.**
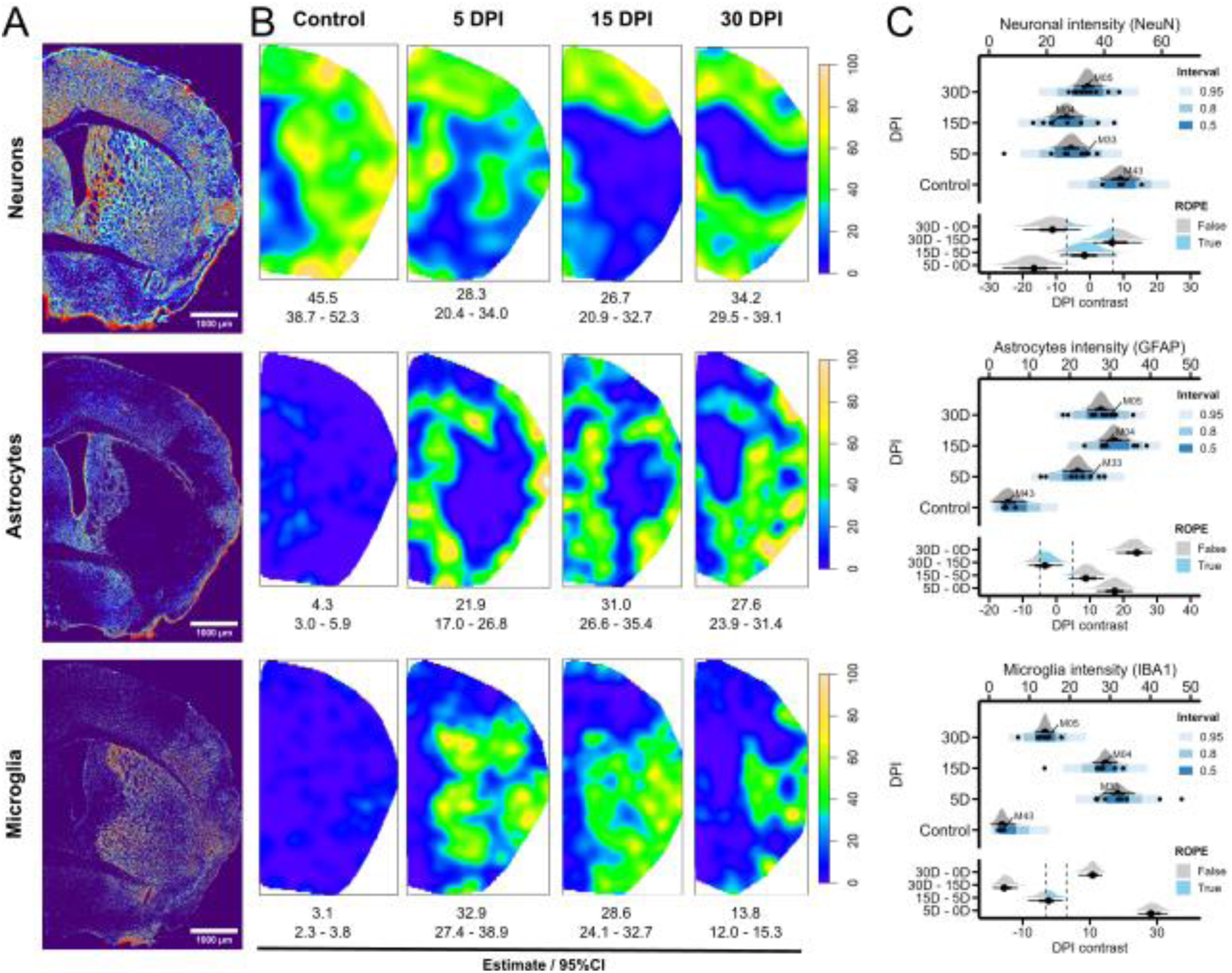
Spatial intensity of neurons, astrocytes, and microglia. **A)** Representative immunolabeling of NeuN (neurons), GFAP (astrocytes), and IBA1 (microglia) at 5 DPI. Brains are displayed using the thermal lookup table (FIJI) to facilitate the visualization of stained cells. **B)** Representative smoothed intensity kernels (density function [sigma = 0.3 (NeuN); sigma = 0.2 (GFAP, IBA1)], spatstat) for NeuN, IBA1, and GFAP at each day post-ischemia (DPI). Mean intensity estimates with 95% credible intervals (CI) are shown below each plot. **C)** Composite graphs showing Bayesian posterior distributions of the spatial intensity for neurons (NeuN, top), astrocytes (GFAP, middle), and microglia (IBA1, bottom). Mean point estimates and their uncertainty are plotted as half-eye and point intervals (stat_halfeye and stat_interval functions, ggdist). Raw data (black dots) is accompanied by prediction intervals (Brewer scale). The contrast between time points of interest (emmeans function [contrast(method = “revpairwise”], emmeans) is displayed at the bottom of each composited graph. The region of practical equivalence (ROPE) encloses the values falling within intragroup variance and, therefore, considered equivalent to null. Probability within the rope is calculated using the hypothesis function from brms. M# labels (unique animal ID) signal the individuals plotted in panels A.

Using PPA, we observed that reactive astrocytes exhibited a peak in spatial intensity at 15 DPI (31, 95% CI = 27.4 - 34.8), which is sustained at later time points (−3.4, 95% CI = −7.6 - 0.77, ROPE = 0.71) (Figure 1B-C, middle). This represents a 6-fold increase compared to the baseline (22.9, 95% CI = 18.3 - 27.6, ROPE = 0). Conversely, the spatial intensity of reactive microglia peaked at 5 DPI (31.9, 95% CI = 28.3 - 36.1) and progressively decreased until 30 DPI (10.5, 95% CI = 7.2 - 13.2) (Figure 1B-C, bottom). We note that those estimates are not directly comparable due to the differying expression of GFAP and IBA1. Furthermore, the negative correlation between the spatial intensity of glia and neurons (QN, Section 2; Suppl. Figure 2D) warrants analysis by PPA.

Altogether, we show that quantifying the spatial intensity enables assessing the dynamic changes in cell populations in response to injury. This approach is compatible with classical estimations of interhemispheric cell ratios (QN, Section 3; Suppl. Figure 3; Suppl. Table 5), showing similar trends during the injury course mainly in the CNU and CTX (QN, section 3; Suppl. Figure 3D).

### Reallocation of glial scar-forming cells following injury

Brain injury involves the activation and reallocation of glial cells to form a scar (Manrique-Castano and ElAli 2021). We investigated this cell redistribution by estimating the spatial intensity of reactive astrocytes and activated microglia as a function of the spatial intensity of neurons (QN, Section 4). We show that GFAP^+^ and IBA1^+^ cells progressively reallocate into regions of low neuronal spatial intensity, as shown by the higher peaks (*rhohat*) at < ∼ 30-40 in Figure 2A.

**Figure 2.**
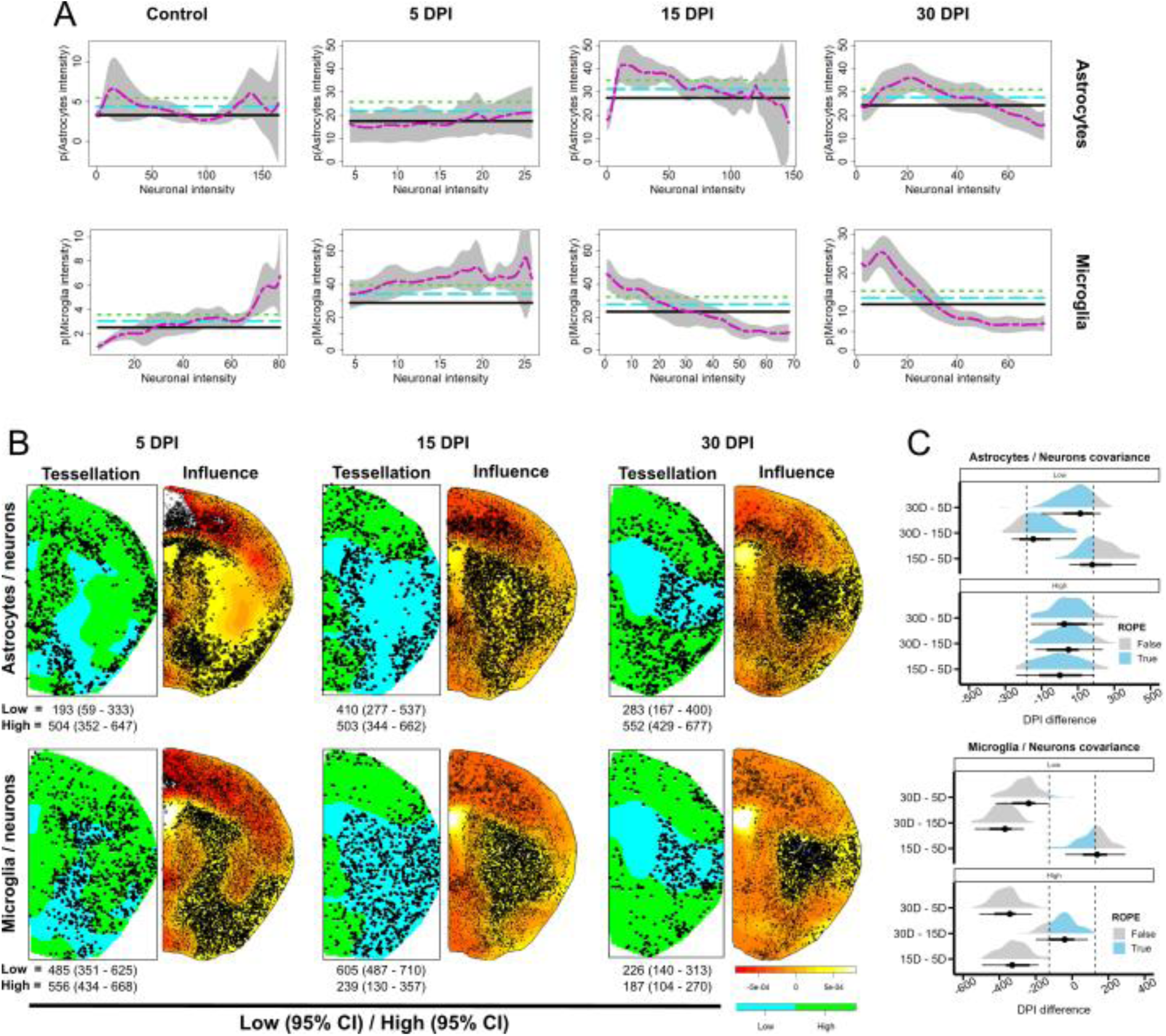
Astrocyte and microglia allocation with respect to neurons. **A)** Conditional intensity distribution estimates (rhohat function, spatstat) for reactive astrocytes (top) and activated microglia (bottom) with respect to the spatial intensity (smoothed density kernel) of neurons. Data are presented as the estimated GFAP or IBA1 intensity (magenta, y-axis) with upper and lower two-sigma confidence intervals (shaded gray area) given the spatial intensity of neurons (x-axis). Horizontal lines represent the mean intensity (dashed line) and the lower (solid black) and upper (dashed green) limits of the two-sigma confidence intervals. Note the different scales, which are weighted by the available ranges of the computed function or constrained for optimal visualization. **B)** (Left, Tessellation) Representative images of tessellated neuronal smooth intensity kernels to segregate regions of low (cyan) and high (green) neuronal spatial intensity (quantiles = 0, 20, 150). The distributions of reactive astrocytes and activated microglia are shown as black dots. Corresponding cell counts in low (top) and high-intensity (bottom) cuadrants and corresponding 95% credible intervals are shown for each DPI. (Right, Influence) Influence (yellow = high, orange = low) of neuronal intensity on glial cell distribution (dfbetas function, spatstat). These maps are generated from superimposed point patterns at each DPI. **C)** Contrast between time points for the posterior distributions of the number of reactive astrocytes and activated microglia in low- and high-intensity neuronal regions (emmeans function [contrast(method = “revpairwise”], emmeans). The mean point estimates and their uncertainty are plotted as half-eye and point intervals (stat_halfeye and stat_interval functions, ggdist). The region of practical equivalence (ROPE) encloses the values falling within intragroup variance and, therefore, considered equivalent to null. Probability within the rope is calculated using the hypothesis function from brms.

Interestingly, the *mppm* [*L_(u)_* = exp (3.1 + 0.002 ± DPI)] for astrocytes shows decreasing slopes from 0.01 at 5 DPI to −0.008 at 30 DPI for time-specific variations (group-level effects) (Suppl. Table 6). The preceding suggests a bimodal dynamic for reactive astrocytes. First, these cells are likely to aggregate in high-intensity (extra-lesional) neuronal regions, shifting later their allocation probability towards low-intensity (intra-lesional) neuronal regions. In contrast, the *mppm* for activated microglia [*L_(u)_* = exp (2.8 + 0.01 ± DPI)] yields negative slopes for all the time-specific variations (Suppl. Table 6). This indicates that IBA1^+^ cells are preferentially positioned in low-intensity (intra-lesional) neuronal regions as the injury progresses.

We also examined the reallocation of reactive glia in neuronal regions using tessellations (quantiles = 0, 20, 150) (QN, Section 5), representing intra-lesional (low spatial intensity) and extra-lesional (high spatial intensity) zones (Figure 2B, Suppl. Table7). Control animals were excluded from this analysis, given the lack of reactive glia labeling in healthy brains.

The results reveal that reactive astrocytes progressively populate low-intensity neuronal regions after the first week post-ischemia (15-5 DPI = 216, 95% CI = 54 - 372, ROPE = 0.35) (Figure 2C, top). Yet, this model yields considerable uncertainty in the estimates, probably due to different infarct sizes and individual evolution of the injury (Suppl. Table 7). Conversely, we predict a persistent distribution of reactive astrocytes in extra-lesional regions throughout injury (Figure 2B-C, top), as implied previously by the *mppm* models. On the other hand, our model predicts that active microglia are increasingly confined to intra-lesional regions (Figure 3B-C, bottom). This entails a ∼77% reduction in IBA1^+^ cells in high-intensity neuronal regions at 30 DPI (30-5 DPI = −370, 95% CI = - 485 - 249, ROPE = 0.005), as well as a reduction in intra-lesional areas (−379, 95% CI = −493 to −260, ROPE = 0.007). This is congruent with the resolution of the inflammatory response. We also found that intra-lesional regions strongly influence glial distribution in the striatum at the level of the MCA territory (Figure 3B; QN, Section 5.8). Notably, the injured area shapes an exclusion zone for GFAP^+^ cells at 5 DPI.

**Figure 3.**
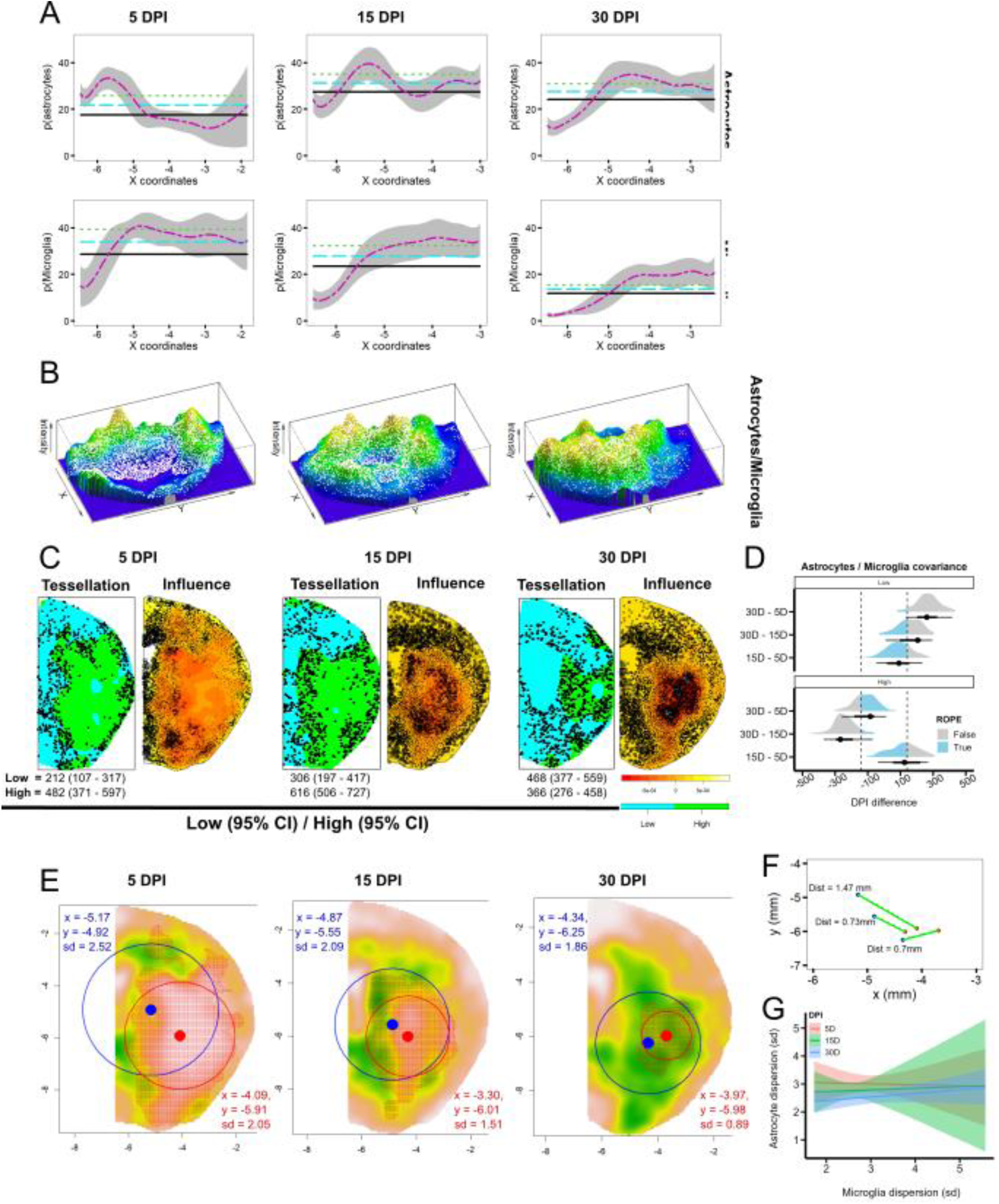
Structuration of a layered glial scar. **A)** Conditional intensity distribution for reactive astrocytes (GFAP^+^) and activated microglia (IBA1^+^) with respect to the x-coordinates of the ischemic hemisphere (rhohat function, spatstat). Data are presented as the estimated GFAP or IBA1 spatial intensity (magenta) with upper and lower two-sigma confidence intervals (shaded gray area). Horizontal lines represent the pooled mean intensity (dashed line) and the lower (solid black) and upper (dashed green) limits of the two-sigma confidence intervals. **B)** Perspective plots (persp function, graphics) of superimposed brains showing the spatial intensity of reactive astrocytes (256 topo colors) and activated microglia (white dots). Reactive astrocytes form a ring that progressively encloses activated microglia in intra-lesional regions. **C)** (Left Tessellation) Representative images of tessellated smooth intensity kernels to segregate regions of low (cyan) and high (green) microglia spatial intensity (quantiles = 0, 20, 150). Astrocytes are displayed as black dots. Corresponding cell counts in low (top) and high (bottom) intensity quadrants for each DPI are shown with 95% credible intervals below each plot. (Right, Influence) Estimated influence (yellow = high, orange = low) of activated microglia on reactive astrocyte distribution (dfbetas function, spatstat). These maps are generated from superimposed point patterns at each DPI. **D)** Contrast between time points for the posterior distributions of the number of reactive astrocytes in low- and high-intensity microglial regions (emmeans function [contrast(method = “revpairwise”], emmeans). The mean point estimates and their uncertainty are plotted as half-eye and point intervals (stat_halfeye and stat_interval functions, ggdist). The region of practical equivalence (ROPE) encloses the values falling within intragroup variance and, therefore, considered equivalent to null. Probability within the rope is calculated using the hypothesis function from brms. **E)** Raster layers (raster function, raster) for reactive astrocytes (terra colors) in superimposed brains. The green color depicts the regions with high cell clustering. Dots and circles depict the centroid and standard deviation for reactive astrocytes (blue) and activated microglia (red). Numerical summaries are displayed in each image with the same color code for both cell types. **F)** Distance between the centroids of high-density raster layers of reactive astrocytes (blue) and activated microglia (red) (5 DPI = 1.47; 15 DPI = 0.73; 30 DPI = 0.7). **G)** Correlation between the point patterns’ standard deviation of reactive astrocytes and activated microglia. Comparable intercepts and slopes (5 DPI, red; 15 DPI, green; 30 DPI, blue) are displayed in the graph. The regression shows a low explained variance (R^2^ = 0.28, 95% CI = 0.05, 0.47).

These results highlight the potential of the relative distribution (rhohat) and cell covariance (tessellations, quadrant counts) estimates to elucidate the complex interactions and spatiotemporal dynamics of different cell types in response to brain injury. This approach can quantitatively characterize the positioning over time of reactive astrocytes and activated microglia relative to neuronal intensity or other spatial features of interest, allowing the modeling of treatments aiming at modulating glial scar properties.

### Astrocytes and microglia form a two-layered glial scar

Next, we examined the interaction between reactive astrocytes and activated microglia at the interface of the glial scar (QN, Section 6). First, we calculated the relative distribution of glial cells with respect to the X-axis (Figure 3A). The graphical results show that reactive astrocytes display a maximum likelihood over the average in extra-lesional areas at 5 DPI. At this stage, we observed a ∼ 2 mm ring of reactive cells around the injury zone, heavily populated by activated microglia (Figure 3B). This ring is narrowed to ∼ 500 µm at 15 DPI, when the mass of reactive astrocytes advances ∼ 200 µm towards the dorsal cortex (Figure 3A). In the later stage, the spatial intensity of reactive astrocytes is prominent in intra-lesional regions between −5 and −3 x-coordinates. Accordingly, the *mppm* [*L_(u)_* = exp (3.2 + 0.007 ± DPI)] (QN, Section 6.1) yielded positive slopes at 15 DPI [exp(0.02)] and 30 DPI [exp (0.17)] (Suppl. Table 8).

On the other hand, activated microglia show a consistent maximum likelihood in intra-lesional areas (∼ -4 - -2 x-coordinates) at all time points (Figure 3A). Besides, the *mppm* shows [*L_(u)_* = exp (4.1 + 0.22 ± DPI)] increasing slopes from exp(− 0.10) at 5DPI to exp (0.15) at 30 DPI (Suppl. Table 8), denoting a progressive centering of the cell mass in intra-lesional regions as shown in Suppl. Figure 4A. The preceding pattern is consistent with the resolution of inflammatory cascades and might imply that astrocytes “sweep” immune cells into intra-lesional areas.

Next, the analysis of GFAP/IBA1 covariance by *mppm* (QN, section 6.3, Suppl. Table 9) suggests a negative correlation (slope, −0.002) that decreases over the injury course (15 DPI = 0.01; 30 DPI = 0.02). This is consistent with the formation of two layers of reactive glial cells that intermingle progressively. Alternatively, analysis using tessellated smoothed density kernels of activated microglia (Figure 3C-D) indicates that reactive astrocytes allocation progressively increases in low-intensity microglial regions by ∼220% from 5 DPI to 30DPI (255, 95% CI = 139-373, ROPE = 0.15) (Suppl. Table 10). Moreover, influence plots (Figure 3C) show that from the second week after injury, astrocyte allocation is strongly influenced by low-density microglial regions, mainly in the cortex.

Taken together, these data support a model where GFAP^+^ and IBA1^+^ cells intermingle progressively during the injury course. Still, astrocyte allocation is prominent in low-intensity microglial areas, consistent with a compartmentalized glial scar with reactive astrocytes (in extra-lesional regions) enclosing activated microglia in an inner layer (in intra-lesional regions).

To further investigate the dynamics of the two glial layers, we analyzed the distance between the centroids of GFAP^+^ and IBA1^+^ cell masses (QN, Section 7). Raw data plots are shown in Suppl. Figure 4B-C. The results show a minimum distance between centroids at 15 DPI (600 μm) (15 - 5 DPI = −0.53, 95% CI = −0.93 to −0.13, ROPE = 0.35) and sustained thereafter (Suppl. Figure 4D). Yet, the astrocyte cell mass shifts its allocation at 30 DPI, as seen in Figure 3E (mean) and Suppl. Figure 4B (single brains). Furthermore, analysis of cell dispersion (QN, Section 7) reveals the higher clustering for reactive astrocytes at 30 DPI and for reactive microglia at 5 DPI (Suppl. Figure 4E). Remarkably, the uncertainty of astrocyte clustering increases as microglial cells disperse (Figure 3G).

The calculation of relative distributions uncovered the progressive mixing of reactive astrocytes and activated microglia at the glial scar. Our results show that reactive astrocytes enclose activated microglia from the second week after injury, and that the microglial distribution in extra-lesional regions strongly influences the boundaries of the reactive astrocyte layer. In addition, raster layers allow an accurate estimation of the dynamics of highly clustered cells that form the glial scar walls.

### Glial scar arrangement in intra-lesional regions

To gain further insights into the structure of the glial scar, we expanded our analysis to intra-lesional regions (QN, Section 8). We defined rectangular ROIs that cover the MCA territory from the ventricular zone to the outer edge of the cerebral cortex (Suppl. Figure 1), and conducted a nonparametric estimation of the conditional distribution of reactive astrocytes with respect to the distance to activated microglia.

The analysis indicates that the spatial intensity of reactive astrocytes is sustained within ∼30 µm of the nearest microglia up to 15 DPI (Figure 4A). However, the peak at > 30 µm at 30 DPI suggests a more distant allocation at this stage. Accordingly, the *mppm* (Suppl. Table 11) yields a higher reactive astrocyte allocation the more distance to activated microglia as the injury progresses [15 DPI = exp(−2.7); 30 DPI = exp(−23.5)]. The preceding suggests a meaningful decoupling of reactive astrocytes and activated microglia when the latter are constrained to intra-lesional regions.

**Figure 4.**
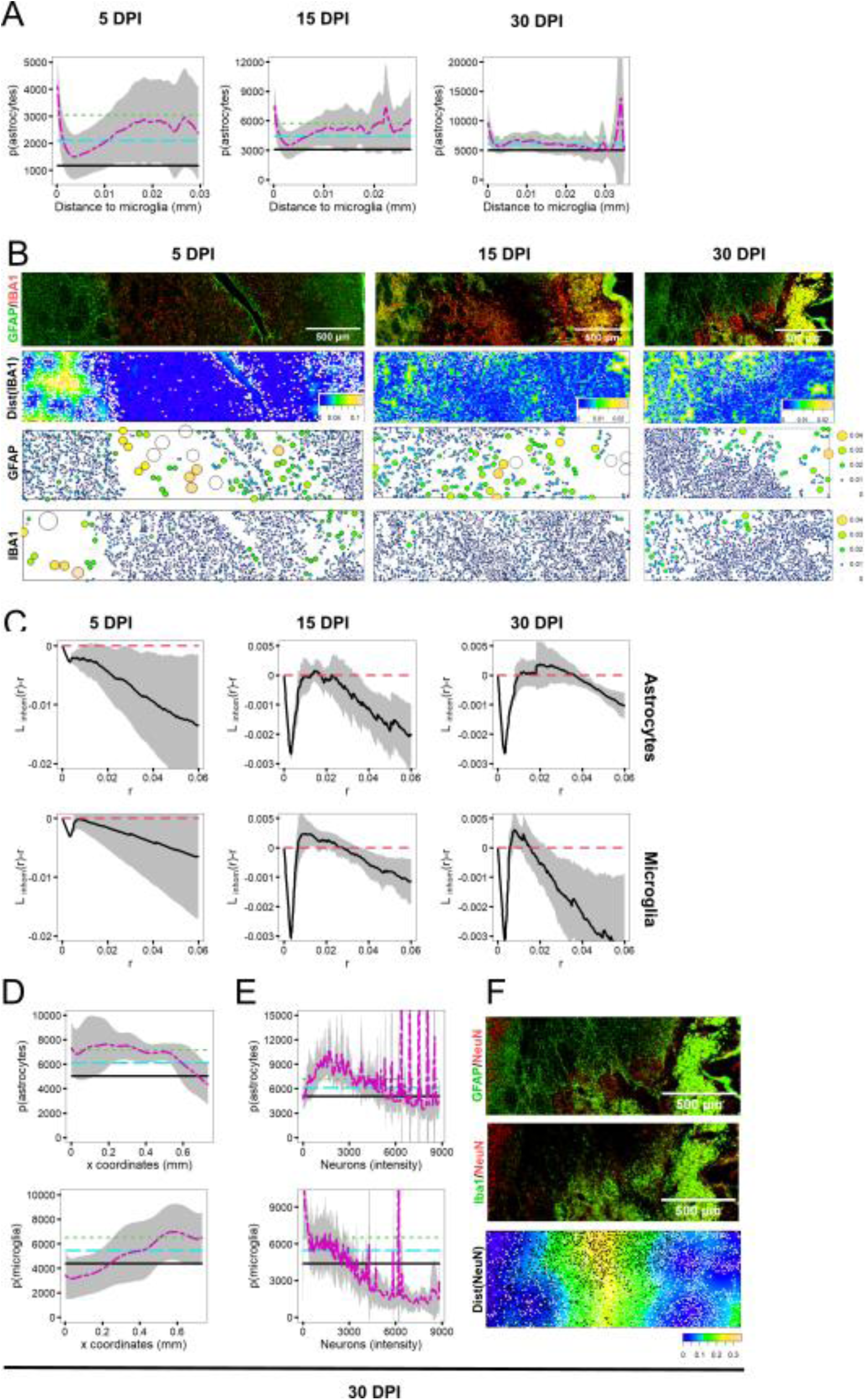
Glial scar in intra-lesional regions. **A)** Conditional intensity distribution (rhohat function, spatstat) for reactive astrocytes with respect to the distance to nearest activated microglia (distfun function, spatstat). Data are presented as the estimated GFAP^+^ cells spatial intensity (magenta) with upper and lower two-sigma confidence intervals (shaded gray area). Horizontal lines represent the pooled mean intensity (dashed line) and the lower (solid black) and upper (dashed green) limits of the two-sigma confidence intervals. **B)** (first row) Representative images of GFAP (green)/IBA1 (red) staining in intra-lesional regions. (second row) distance maps (distfun function, spatstat) of activated microglia in the same region (256 topo colors). Green and yellow zones depict the areas where the distance between activated microglia (low spatial intensity) is more extensive. Reactive astrocytes are represented as white dots. (third and fourth rows) Stienen maps/diagrams (stienen function, spatstat) of GFAP^+^ and IBA1^+^ reactive glia depicting circles around each data point with a diameter corresponding to the distance to the nearest neighbor. **C)** Inhomogeneous L-function (Linhom function, spatstat) to estimate long-scale cumulative distributions of reactive astrocytes and activated microglia. The dashed red line is centered at 0 and corresponds to a regular point pattern. Over and below the line signal clustering and inhibition (independent), respectively. The estimates are based on Ripley’s isotropic correction (black line) with upper and lower limits of two-sigma CI (shadowed region). **D-E)** Conditional spatial intensity distribution for reactive astrocytes and active microglia with respect to the x-axis of indicated ROIs (rhohat function, spatstat), and neuronal spatial intensity in intra-lesional regions, respectively. Data are presented as the estimated GFAP^+^ cells spatial intensity (magenta) with upper and lower two-sigma confidence intervals (shaded gray area). Horizontal lines represent the pooled mean intensity (dashed line) and the lower (solid black) and upper (dashed green) limits of the two-sigma confidence intervals. **F)** Representative images of GFAP (green)/ NeuN (red) (first row) and IBA1 (green)/ NeuN (red) (second row) staining in intra-lesional regions demarking the allocation of glial cells with respect to neurons. (Third row) Distance function estimate (distfun function, spatstat) for neurons in intra-lesional areas. Green and yellow zones depict the areas where the distance between neurons (low spatial instensity) is larger.

We also analyzed the interaction within cells (QN, section 9). The results denote that, on the long scale, reactive astrocytes transition from an independent to a regular distribution, becoming more evenly spaced over time (QN, section 9.1; Figure 4C, top). In contrast, activated microglia are likely to maintain an independent distribution in the chronic phase of the injury and show less variability in their spatial distribution (QN, section 9.2; Figure 4C, bottom). Otherwise, on the short scale, activated microglia exhibit higher clustering than astrocytes at 15 DPI and display a more restricted behavior between ∼20 and ∼40 µm at 30 DPI (Figure 4C). This implies that the distribution of reactive astrocytes is more dynamic than that of activated microglia, possibly because their allocation at the edges of the injured tissue.

In addition, we analyzed the distribution of reactive astrocytes and activated microglia with respect to the x-coordinates of the ROI at 30 DPI. Our models indicate that reactive astrocytes are concentrated in the innermost part of the ischemic hemisphere [*L_(u)_* = exp (9.1 - 0.9)], while activated microglia are located near the hemispheric boundary [*L_(u)_* = exp (8.4 - 0.3)] (Figure 4D). In the MCA territory, both cell types are prominently positioned in areas with low neuronal intensity (QN, Section 8.3; see Figure 4E). These findings support the notion of a compartmentalized glial scar in which the outer layer of reactive astrocytes pushes tissue debris full of activated microglia to the borders of the brain.

### TDA supports the spatiotemporal dynamics of reactive glia observed in PPA

TDA revealed a correlation in the density of reactive astrocytes and neurons (p < 0.001) during the first two weeks post-ischemia. Conversely, TDA tracked a shift of activated microglia from even distribution in control conditions to clustering in specific (intra-lesional) regions after injury (p < 0.01). Interestingly, the distribution of activated microglia appears to be correlated with the high-density neuronal regions where reactive astrocytes initially aggregated. Analysis of Betti curves obtained from TDA of NeuN^+^, GFAP^+,^ and IBA1^+^ cells positions and pairwise combinations of cell types revealed that reactive astrocytes and activated microglia are present in regions of low neuronal density (Figure 5A).

**Figure 5.**
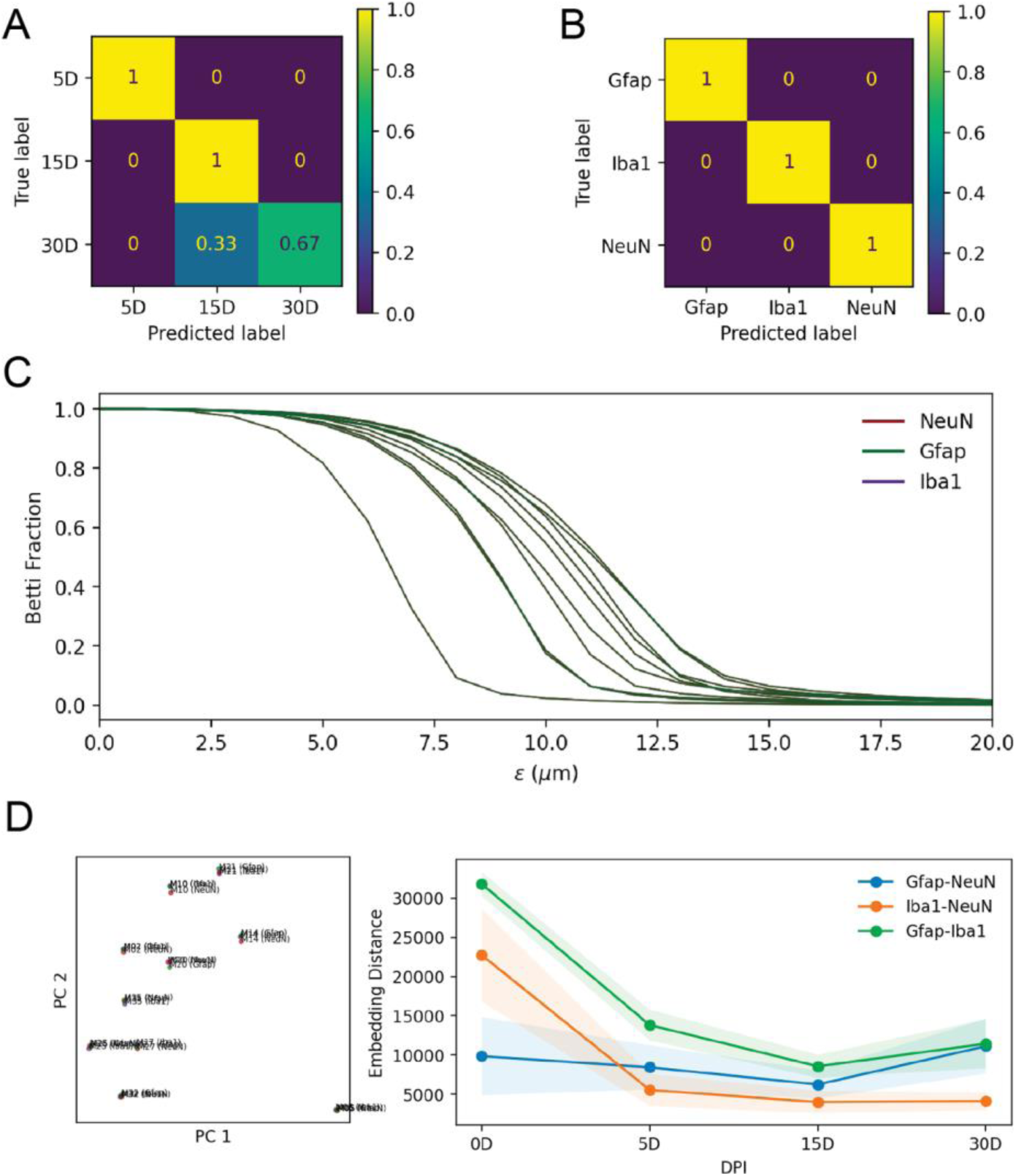
Recovery of DPI and cell type from topological features. **A)** Confusion matrix for the prediction of days post-injury with an overall accuracy of 77.8% using dimension 0 and dimension 1 topological features. **B)** Confusion matrix for the prediction of cell type with 100% accuracy using dimension 0 and dimension 1 topological features. **C)** Betti curves are clustered by mouse identity. **D)** Principal component analysis of Betti curves reveals clustering by mouse identity. Jitter added to plot for visual clarity. Embedding distance between cell types over time reveals co-occurrence of astrocytes (Gfap) and neurons (NeuN) during the first 5-15 days before shifting to low neuronal density regions at 30 DPI.

### TDA predicts DPI using the position of single-cell types

We used a topological machine learning-based approach to predict DPI based on cell positions. We first applied persistent homology analysis to the point clouds to obtain topological features of the data. Subsequently, we predicted DPI using SVR with RBF kernel. The results show that this approach can accurately predict DPI based on the positions (point clouds) of neurons, reactive astrocytes, and activated microglia, with a mean absolute error (MAE) of 0.84 days and a root mean squared error (RMSE) of 1.16 days (Suppl. Table 12).

To further validate the effectiveness of this model, we compared it with two baseline methods: a linear regression model and a neural network model with two hidden layers. The linear regression model achieved a higher MAE of 1.58 days and a higher RMSE of 2.05 days, while the neural network model achieved a similar MAE of 0.88 days but a higher RMSE of 1.39 days (Suppl. Table 13). The preceding denotes that our topological machine learning model outperforms the baseline methods regarding prediction accuracy.

We also performed a feature importance analysis to identify the most informative features for predicting DPI. Our analysis revealed that the positions of activated microglia were the most informative features, followed by the positions of neurons and reactive astrocytes (Suppl. Table 14).

Overall, our results demonstrate the effectiveness of our topological machine learning-based approach for predicting the post-ischemia time point using point cloud data of neurons, astrocytes, and microglia positions. This allows the use of the topological features of glial cell distribution to inform about injury progression in CNS diseases.

## Discussion

Traditional methods to characterize the heterogeneous response of glia, based on cell counting or quantification of glial markers stained area (Anderson et al. 2016; Okada et al. 2006; Buscemi et al. 2019), do not fully capture the complex dynamics of glial scars. In this study, we developed PPA/TDA-based approaches to investigate the structure and spatiotemporal arrangement of glial scars in the mouse brain following injury. With this approximation, researchers can uncover distribution patterns of reactive glia at different scales. The use of the *rhohat* and *ppm* frameworks from the *spatstat* R-package enables the assessment of the spatial distribution of reactive astrocytes and activated microglia, taking into consideration cell-to-cell interactions and the heterogeneity of the spatial process (*Spatial Point Patterns: Methodology and Applications with R* 2015). To our knowledge, the use point pattern-based approaches in biomedicine is restricted to cancer biology (Maisel et al. 2022; Kaufmann et al. 2021).

Numerous approaches exist that demonstrate the impact of microglia (Zhang et al. 2023; Bellver-Landete et al. 2019; Fu et al. 2020), astrocytes (Anderson et al. 2016; Gu et al. 2019; Herrmann et al. 2008), or NG2 (Rodriguez et al. 2014; Hesp et al. 2017) dysregulation/ablation in scar formation and their subsequent effects on neurological recovery. The assessment of these and other therapeutic approaches could benefit from analyzing the fine traits of cell reallocation, covariance, and interaction. Indeed, PPA or TDA-based approaches can benefit the assessment of cell or protein distribution in other CNS diseases such as Alzheimer’s (Spangenberg et al. 2019; Al-Onaizi et al. 2022), Huntington’s (Crapser et al. 2019), or Parkinson’s disease (Sekiya et al. 2022).

(Wanner et al. 2013) demonstrated that glial scar borders are constituted by proliferating astrocytes, which serve to contain fibrotic and inflammatory cells. However, their findings do not incorporate a quantitative model that facilitates predictions regarding cellular covariance or temporal evolution—capabilities inherent to our approach. Our model elucidates the distribution patterns of reactive astrocytes and activated microglia, thereby enabling the exploration of alternative hypotheses regarding fibrotic scar deposition (Fernández-Klett and Priller 2014). Although astrocytes are commonly believed to be primary contributors to the formation of fibrotic scar through the production of extracellular matrix (ECM) proteins like tenascin-C (TnC) (Dzyubenko et al. 2018; Dzyubenko et al. 2022), their predominant localization in regions outside the lesion suggests a potential key role for other cell types. Specifically, pericytes/PDGFR- β+ cells might play a central role in this process, as indicated by other studies (Dias et al. 2021; Soderblom et al. 2013; Göritz et al. 2011) .

Recently, TDA analysis has gained attention in oncology as a tool to determine cell architecture, disease classification, computer-aided diagnosis, prediction of treatment response (Bukkuri, Andor, and Darcy 2021; Singh et al. 2023), or to explore the gene regulatory network (Masoomy et al. 2021). Also, TDA has been used to quantify cell-to-cell interactions (Bhaskar, Zhang, and Wong 2021), track their collective motions (Bonilla, Carpio, and Trenado 2020), or feature emerging traits in cell dynamics (Dawson et al. 2022). In neurobiology, TDA applications are limited to EGG (Yamanashi et al. 2021) or fMRI analysis (Saggar et al. 2022) for the analysis of brain networks (Das, Anand, and Chung 2023). Here, we show the first application of TDA to analyze the topological arrangement of nervous cells (point clouds) extracted from widefield microscopy images. This entails a step towards an unbiased and reproducible analysis of cell distribution in fixed brain tissue, which can be extrapolated to other body organs.

Automated/unbiased PPA for analyzing glial scars or cell redistribution has some limitations. First, the accuracy of the initial cell detection (performed in QuPath) can be influenced by several factors, including immunolabeling efficiency, cell size, and cell overlapping. In particular, we faced difficulty segmenting individual GFAP^+^ and IBA1^+^ cells. Conversely, using nuclear markers like NeuN is advantageous for segmentation algorithms like watershed. Several algorithms are suitable for segmenting quasi-circular or oval cells (Al-Kofahi et al. 2018; Greenwald et al. 2021; Stringer et al. 2020). However, the segmentation of highly branched and overlapping cells such as reactive astrocytes and activated microglia is a challenge not yet addressed. Secondly, the estimation of spatial intensity depends on the size and shape of the observation window. The window size can influence the detection of clustering patterns and density calculation, so it should be chosen carefully. Third, interpreting *ppm* and *mppm* models requires careful consideration of the assumptions and parameters. Unfortunately, there are still no tools in *spatstat* for *mppm* diagnostics or goodness-of-fit. For this reason, we complete our analysis with Bayesian models. Furthermore, running *ppm* models to large datasets can be computationally intensive and time-consuming, which may limit its scalability for some applications involving many cells. Finally, the interpretation of spatial intensity maps can be influenced by choice of color scales and contour lines, which emphasize the importance of proper modeling.

Our study shows that the limitations of 2D PPA for the study of reactive glia and scar formation in the CNS are surprassed by 3D TDA, which captures the full complexity of the 3D microenvironment in which cells interact and migrate. Still, TDA supported the results of PPA, indicating that astrocytes initially aggregate in high-density neuronal regions during the first week post-injury, before shifting to low-density neuronal regions in the second week.

TDA, particularly through the use of persistent homology, relies on the creation of simplicial complexes from the data. In the presence of noise, these complexes can capture spurious topological features that do not reflect the underlying geometry or structure of the data. This can lead to misinterpretations. Moreover, the scale at which the data is analyzed plays a crucial role; small-scale features might be missed, while large-scale features might overshadow more intricate details. Also, constructing simplicial complexes, computing homology groups, and tracking persistence across scales can be computationally demanding. As the size and dimensionality of the data increase, the computational cost associated with TDA can escalate rapidly. This can limit the real-time or on-the-fly applications of TDA for big datasets. To overcome these limitations, we used a sampling strategy to subset our data.

On the other hand, Bayesian modeling with parameter estimation and uncertainty quantification offers several advantages of traditional hypothesis testing (Kruschke and Liddell 2018). This framework enables the direct tunning of model parameters and obtaining transparent contrast and effect sizes based on the full posterior distribution. We found the *brms* and *emmeans* R-packages fundamental to contrasts grouping variables of interest, and we promote its use for proper statistical inference based on estimation and uncertainty, which is particularly relevant for small sample sizes such as those typical of biomedicine.

In conclusion, our study has successfully applied PPA and TDA to investigate the spatial distribution and evolution of neurons, astrocytes, and microglia in the injured brain after ischemic stroke. Also, it demonstrated the power and versatility of machine learning techniques to predict days post-injury from cell positions with high accuracy. This result not only validates the robustness of TDA for analyzing complex biological data, but also highlights its potential for aiding in the development of novel diagnostic and therapeutic strategies for neurological disorders and injuries. Our findings contribute to a better understanding of the dynamic cellular interactions occurring in the brain following ischemic stroke and provide a foundation for future research.

## Acknowledgments

D.M-C is funded by Fonds de recherche du Québec (FRQS)-Santé (case - 318466). D.B. is funded by a Yale-Boehringer Ingelheim Biomedical Data Science Fellowship.

## Code availability

Raw data and R/Julia code for full reproducibility and reusability of our approach are available in the OSF (DOI 10.17605/OSF.IO/3VG8J), Zenodo (https://doi.org/10.5281/zenodo.8399976) and GitHub (https://github.com/elalilab/Stroke_GlialScar_PPA-TDA) repositories. The maintenance is performed by the Laboratory of Neurovascular Interactions (Université Laval), directed by Ayman Eali.

## References

Bradbury, Elizabeth J., and Emily R. Burnside. 2019. “Moving beyond the Glial Scar for Spinal Cord Repair”. Nature Communications 10 (1). 10.1038/s41467-019-11707-7.

Wanner, I. B., M. A. Anderson, B. Song, J. Levine, A. Fernandez, Z. Gray-Thompson, Y. Ao, and M. V. Sofroniew. 2013. “Glial Scar Borders Are Formed by Newly Proliferated Elongated Astrocytes That Interact to Corral Inflammatory and Fibrotic Cells via STAT3-Dependent Mechanisms after Spinal Cord Injury”. Journal of Neuroscience 33 (31): 12870–86. 10.1523/jneurosci.2121-13.2013.

Hackett, Amber R., and Jae K. Lee. 2016. “Understanding the NG2 Glial Scar after Spinal Cord Injury”. Frontiers in Neurology 7 (November). 10.3389/fneur.2016.00199.

Bellver-Landete, Victor, Floriane Bretheau, Benoit Mailhot, Nicolas Vallières, Martine Lessard, Marie-Eve Janelle, Nathalie Vernoux, et al. 2019. “Microglia Are an Essential Component of the Neuroprotective Scar That Forms after Spinal Cord Injury”. Nature Communications 10 (1). 10.1038/s41467-019-08446-0.

Zhang, C, J Kang, X Zhang, Y Zhang, N Huang, and B Ning. 2022. “Spatiotemporal Dynamics of the Cellular Components Involved in Glial Scar Formation Following Spinal Cord Injury.”. Biomed Pharmacother 153: 113500.

Conforti, P, S Mezey, S Nath, YH Chu, SC Malik, Santamaría JC Martínez, SS Deshpande, L Pous, B Zieger, and C Schachtrup. 2022. “Fibrinogen Regulates Lesion Border-Forming Reactive Astrocyte Properties after Vascular Damage.”. Glia 70: 1251– 66.

Voskuhl, R. R., R. S. Peterson, B. Song, Y. Ao, L. B. J. Morales, S. Tiwari-Woodruff, and M. V. Sofroniew. 2009. “Reactive Astrocytes Form Scar-Like Perivascular Barriers to Leukocytes during Adaptive Immune Inflammation of the CNS”. Journal of Neuroscience 29 (37): 11511–22. 10.1523/jneurosci.1514-09.2009.

Yoshizaki, S, T Tamaru, M Hara, K Kijima, M Tanaka, DJ Konno, Y Matsumoto, Y Nakashima, and S Okada. 2021. “Microglial Inflammation after Chronic Spinal Cord Injury Is Enhanced by Reactive Astrocytes via the Fibronectin/β1 Integrin Pathway.”. J Neuroinflammation 18: 12.

Anderson, Mark A., Joshua E. Burda, Yilong Ren, Yan Ao, Timothy M. O’Shea, Riki Kawaguchi, Giovanni Coppola, Baljit S. Khakh, Timothy J. Deming, and Michael V. Sofroniew. 2016. “Astrocyte Scar Formation Aids Central Nervous System Axon Regeneration”. Nature 532 (7598): 195–200. 10.1038/nature17623.

Fu, H, Y Zhao, D Hu, S Wang, T Yu, and L Zhang. 2020. “Depletion of Microglia Exacerbates Injury and Impairs Function Recovery after Spinal Cord Injury in Mice.”. Cell Death Dis 11: 528.

Yang, Tuo, YuJuan Dai, Gang Chen, and ShuSen Cui. 2020. “Dissecting the Dual Role of the Glial Scar and Scar-Forming Astrocytes in Spinal Cord Injury”. Frontiers in Cellular Neuroscience 14 (April). 10.3389/fncel.2020.00078.

Kamphuis, W, L Kooijman, M Orre, O Stassen, M Pekny, and EM Hol. 2015. “GFAP and Vimentin Deficiency Alters Gene Expression in Astrocytes and Microglia in Wild-Type Mice and Changes the Transcriptional Response of Reactive Glia in Mouse Model for Alzheimer’s Disease.”. Glia 63: 1036–56.

Schacke, S, J Kirkpatrick, A Stocksdale, R Bauer, C Hagel, LB Riecken, and H Morrison. 2022. “Ezrin Deficiency Triggers Glial Fibrillary Acidic Protein Upregulation and a Distinct Reactive Astrocyte Phenotype.”. Glia 70: 2309–29.

Lu, X, F Lu, J Yu, X Xue, H Jiang, L Jiang, and Y Yang. 2021. “Gramine Promotes Functional Recovery after Spinal Cord Injury via Ameliorating Microglia Activation.”. J Cell Mol Med 25: 7980–92.

Buscemi, Lara, Melanie Price, Paola Bezzi, and Lorenz Hirt. 2019. “Spatio-Temporal Overview of Neuroinflammation in an Experimental Mouse Stroke Model”. Scientific Reports 9 (1). 10.1038/s41598-018-36598-4.

Manrique-Castano, Daniel, Egor Dzyubenko, Mina Borbor, Paraskevi Vasileiadou, Christoph Kleinschnitz, Lars Roll, Andreas Faissner, and Dirk M. Hermann. 2021. “Tenascin-C Preserves Microglia Surveillance and Restricts Leukocyte and More Specifically, T Cell Infiltration of the Ischemic Brain”. Brain, Behavior, and Immunity 91 (January): 639–48. 10.1016/j.bbi.2020.10.016.

Ito, D, K Tanaka, S Suzuki, T Dembo, and Y Fukuuchi. 2001. “Enhanced Expression of Iba1, Ionized Calcium-Binding Adapter Molecule 1, after Transient Focal Cerebral Ischemia in Rat Brain.”. Stroke 32: 1208–15.

Parra, Edwin Roger. 2021. “Methods to Determine and Analyze the Cellular Spatial Distribution Extracted From Multiplex Immunofluorescence Data to Understand the Tumor Microenvironment”. Frontiers in Molecular Biosciences 8 (June). 10.3389/fmolb.2021.668340.

Jafari-Mamaghani, M, M Andersson, and P Krieger. 2010. “Spatial Point Pattern Analysis of Neurons Using Ripley’s K-Function in 3D.”. Front Neuroinform 4: 9.

Davis, Benjamin M., Manual Salinas-Navarro, M. Francesca Cordeiro, Lieve Moons, and Lies De Groef. 2017. “Characterizing Microglia Activation: a Spatial Statistics Approach to Maximize Information Extraction”. Scientific Reports 7 (1). 10.1038/s41598-017-01747-8.

Prodanov, D, N Nagelkerke, and E Marani. 2007. “Spatial Clustering Analysis in Neuroanatomy: Applications of Different Approaches to Motor Nerve Fiber Distribution.”. J Neurosci Methods 160: 93–108.

Bhaskar, D, WY Zhang, and IY Wong. 2021. “Topological Data Analysis of Collective and Individual Epithelial Cells Using Persistent Homology of Loops.”. Soft Matter 17: 4653–64.

Masoomy, H, B Askari, S Tajik, AK Rizi, and GR Jafari. 2021. “Topological Analysis of Interaction Patterns in Cancer-Specific Gene Regulatory Network: Persistent Homology Approach.”. Sci Rep 11: 16414.

Bonilla, LL, A Carpio, and C Trenado. 2020. “Tracking Collective Cell Motion by Topological Data Analysis.”. PLoS Comput Biol 16: e1008407.

Malavasi, Marco, Manuele Bazzichetto, Stefania Bagella, Vojtěch Barták, Anna Depalmas, Antonello Gregorini, Marta Gaia Sperandii, Alicia T. R. Acosta, and Simonetta Bagella. 2023. “Ecology Meets Archaeology: Past Present and Future Vegetation-Derived Ecosystems Services from the Nuragic Sardinia (1700580 $\Less$Scp$\Greater$BCE$\Less$/Scp$\Greater$)”. People and Nature, March. 10.1002/pan3.10461.

Scarpone, Christopher, Sebastian T. Brinkmann, Tim Große, Daniel Sonnenwald, Martin Fuchs, and Blake Byron Walker. 2020. “A Multimethod Approach for County-Scale Geospatial Analysis of Emerging Infectious Diseases: a Cross-Sectional Case Study of COVID-19 Incidence in Germany”. International Journal of Health Geographics 19 (1). 10.1186/s12942-020-00225-1.

Maisel, Brenton A., Misung Yi, Amy R. Peck, Yunguang Sun, Jeffrey A. Hooke, Albert J. Kovatich, Craig D. Shriver, et al. 2022. “Spatial Metrics of Interaction between CD163-Positive Macrophages and Cancer Cells and Progression-Free Survival in Chemo-Treated Breast Cancer”. Cancers 14 (2): 308. 10.3390/cancers14020308.

Kaufmann, J, CAN Biscio, P Bankhead, S Zimmer, H Schmidberger, E Rubak, and A Mayer. 2021. “Using the R Package Spatstat to Assess Inhibitory Effects of Microregional Hypoxia on the Infiltration of Cancers of the Head and Neck Region by Cytotoxic T Lymphocytes.”. Cancers (Basel) 13.

Vipond, Oliver, Joshua A. Bull, Philip S. Macklin, Ulrike Tillmann, Christopher W. Pugh, Helen M. Byrne, and Heather A. Harrington. 2021. “Multiparameter Persistent Homology Landscapes Identify Immune Cell Spatial Patterns in Tumors”. Proceedings of the National Academy of Sciences 118 (41). 10.1073/pnas.2102166118.

Lawson, Peter, Andrew B. Sholl, J. Quincy Brown, Brittany Terese Fasy, and Carola Wenk. 2019. “Persistent Homology for the Quantitative Evaluation of Architectural Features in Prostate Cancer Histology”. Scientific Reports 9 (1). 10.1038/s41598-018-36798-y.

Amézquita, Erik J., Michelle Y. Quigley, Tim Ophelders, Elizabeth Munch, and Daniel H. Chitwood. 2020. “The Shape of Things to Come: Topological Data Analysis and Biology from Molecules to Organisms”. Developmental Dynamics 249 (7): 816–33. 10.1002/dvdy.175.

Townsend, Jacob, Cassie Putman Micucci, John H. Hymel, Vasileios Maroulas, and Konstantinos D. Vogiatzis. 2020. “Representation of Molecular Structures with Persistent Homology for Machine Learning Applications in Chemistry”. Nature Communications 11 (1). 10.1038/s41467-020-17035-5.

Kanari, Lida, Paweł Dłotko, Martina Scolamiero, Ran Levi, Julian Shillcock, Kathryn Hess, and Henry Markram. 2017. “A Topological Representation of Branching Neuronal Morphologies”. Neuroinformatics 16 (1): 3–13. 10.1007/s12021-017-9341-1.

Nardini, John T., Bernadette J. Stolz, Kevin B. Flores, Heather A. Harrington, and Helen M. Byrne. 2021. “Topological Data Analysis Distinguishes Parameter Regimes in the Anderson-Chaplain Model of Angiogenesis”. Edited by Mark Chaplain. PLOS Computational Biology 17 (6): e1009094. 10.1371/journal.pcbi.1009094.

Dittmar, Michael S., Bijan Vatankhah, Nando P. Fehm, Gerald Retzl, Gerhard Schuierer, Ulrich Bogdahn, Felix Schlachetzki, and Markus Horn. 2005. “The Role of ECA Transection in the Development of Masticatory Lesions in the MCAO Filament Model”. Experimental Neurology 195 (2): 372–78. 10.1016/j.expneurol.2005.05.013.

Schindelin, Johannes, Ignacio Arganda-Carreras, Erwin Frise, Verena Kaynig, Mark Longair, Tobias Pietzsch, Stephan Preibisch, et al. 2012. “Fiji: an Open-Source Platform for Biological-Image Analysis”. Nature Methods 9 (7): 676–82. 10.1038/nmeth.2019.

Bankhead, Peter, Maurice B. Loughrey, José A. Fernández, Yvonne Dombrowski, Darragh G. McArt, Philip D. Dunne, Stephen McQuaid, et al. 2017. “QuPath: Open Source Software for Digital Pathology Image Analysis”. Scientific Reports 7 (1). 10.1038/s41598-017-17204-5.

Baddeley, Adrian, Ege Rubak, and Rolf Turner. 2015. Spatial Point Patterns. Chapman & Hall/CRC Interdisciplinary Statistics. Oakville, MO: Apple Academic Press.

Hijmans, Robert J. 2023. Raster: Geographic Data Analysis and Modeling. https://CRAN.R-project.org/package=raster.

Zhang, Simon, Mengbai Xiao, and Hao Wang. 2020. “GPU-Accelerated Computation of Vietoris-Rips Persistence Barcodes”. In 36th International Symposium on Computational Geometry (SoCG 2020). Schloss Dagstuhl-Leibniz-Zentrum für Informatik.

Hensel, Felix, Michael Moor, and Bastian Rieck. 2021. “A Survey of Topological Machine Learning Methods”. Frontiers in Artificial Intelligence 4 (May). 10.3389/frai.2021.681108.

Ben-Hur, Asa, Cheng Soon Ong, Sören Sonnenburg, Bernhard Schölkopf, and Gunnar Rätsch. 2008. “Support Vector Machines and Kernels for Computational Biology”. Edited by Fran Lewitter. PLoS Computational Biology 4 (10): e1000173. 10.1371/journal.pcbi.1000173.

Dutschmann, Thomas-Martin, Lennart Kinzel, Antonius ter Laak, and Knut Baumann. 2023. “Large-Scale Evaluation of k-Fold Cross-Validation Ensembles for Uncertainty Estimation”. Journal of Cheminformatics 15 (1). 10.1186/s13321-023-00709-9.

Jung, Yoonsuh. 2017. “Multiple Predicting$\Less$i$\Greater$K$\Less$/i$\Greater$-Fold Cross-Validation for Model Selection”. Journal of Nonparametric Statistics 30 (1): 197–215. 10.1080/10485252.2017.1404598.

Bürkner, Paul-Christian. 2018. “Advanced Bayesian Multilevel Modeling with the R Package Brms”. The R Journal 10 (1): 395. 10.32614/rj-2018-017.

Baddeley, Adrian, and Rolf Turner. 2005. “$\Less$b$\Greater$Spatstat$\Less$/b$\Greater$: An$\Less$i$\Greater$R$\Less$/i$\Greater$Package for Analyzing Spatial Point Patterns”. Journal of Statistical Software 12 (6). 10.18637/jss.v012.i06.

Hespanhol, L, CS Vallio, LM Costa, and BT Saragiotto. 2019. “Understanding and Interpreting Confidence and Credible Intervals around Effect Estimates.”. Braz J Phys Ther 23: 290–301.

Hahn, Ute. 2012. “A Studentized Permutation Test for the Comparison of Spatial Point Patterns”. Journal of the American Statistical Association 107 (498): 754–64. 10.1080/01621459.2012.688463.

Sofroniew, Michael V., and Harry V. Vinters. 2009. “Astrocytes: Biology and Pathology”. Acta Neuropathologica 119 (1): 7–35. 10.1007/s00401-009-0619-8.

Unal-Cevik, I, M Kilinç, Y Gürsoy-Ozdemir, G Gurer, and T Dalkara. 2004. “Loss of NeuN Immunoreactivity after Cerebral Ischemia Does Not Indicate Neuronal Cell Loss: a Cautionary Note.”. Brain Res 1015: 169–74.

Manrique-Castano, Daniel, and Ayman ElAli. 2021. “Neurovascular Reactivity in Tissue Scarring Following Cerebral Ischemia”. In Cerebral Ischemia, 111–30. Exon Publications. 10.36255/exonpublications.cerebralischemia.2021.neurovascularreactivit y.

Okada, Seiji, Masaya Nakamura, Hiroyuki Katoh, Tamaki Miyao, Takuya Shimazaki, Ken Ishii, Junichi Yamane, et al. 2006. “Conditional Ablation of Stat3 or Socs3 Discloses a Dual Role for Reactive Astrocytes after Spinal Cord Injury”. Nature Medicine 12 (7): 829–34. 10.1038/nm1425.

2015. Chapman and Hall/CRC Press.

Zhang, Hui, Zhi-Lai Zhou, Huan Xie, Xiao-Bo Tian, Hua-Li Xu, Wei Li, and Shun Yao. 2023. “Microglial Depletion Impairs Glial Scar Formation and Aggravates Inflammation Partly by Inhibiting STAT3 Phosphorylation in Astrocytes after Spinal Cord Injury”. Neural Regeneration Research 18 (6): 1325. 10.4103/1673-5374.357912.

Fu, Haitao, Yanpeng Zhao, Die Hu, Song Wang, Tengbo Yu, and Licheng Zhang. 2020. “Depletion of Microglia Exacerbates Injury and Impairs Function Recovery after Spinal Cord Injury in Mice”. Cell Death &Amp$\Mathsemicolon$ Disease 11 (7). 10.1038/s41419-020-2733-4.

Gu, Yakun, Xueyan Cheng, Xiao Huang, Yimin Yuan, Shangyao Qin, Zijian Tan, Dan Wang, Xin Hu, Cheng He, and Zhida Su. 2019. “Conditional Ablation of Reactive Astrocytes to Dissect Their Roles in Spinal Cord Injury and Repair”. Brain Behavior, and Immunity 80 (August): 394–405. 10.1016/j.bbi.2019.04.016.

Herrmann, Julia E., Tetsuya Imura, Bingbing Song, Jingwei Qi, Yan Ao, Thu K. Nguyen, Rose A. Korsak, Kiyoshi Takeda, Shizuo Akira, and Michael V. Sofroniew. 2008. “STAT3 Is a Critical Regulator of Astrogliosis and Scar Formation after Spinal Cord Injury”. The Journal of Neuroscience 28 (28): 7231–43. 10.1523/jneurosci.1709-08.2008.

Rodriguez, J. P., M. Coulter, J. Miotke, R. L. Meyer, K.-I. Takemaru, and J. M. Levine. 2014. “Abrogation of -Catenin Signaling in Oligodendrocyte Precursor Cells Reduces Glial Scarring and Promotes Axon Regeneration after CNS Injury”. Journal of Neuroscience 34 (31): 10285–97. 10.1523/jneurosci.4915-13.2014.

Hesp, Zoe C., Rim Y. Yoseph, Ryusuke Suzuki, Peter Jukkola, Claire Wilson, Akiko Nishiyama, and Dana M. McTigue. 2017. “Proliferating NG2-Cell-Dependent Angiogenesis and Scar Formation Alter Axon Growth and Functional Recovery After Spinal Cord Injury in Mice”. The Journal of Neuroscience 38 (6): 1366–82. 10.1523/jneurosci.3953-16.2017.

Spangenberg, Elizabeth, Paul L. Severson, Lindsay A. Hohsfield, Joshua Crapser, Jiazhong Zhang, Elizabeth A. Burton, Ying Zhang, et al. 2019. “Sustained Microglial Depletion with CSF1R Inhibitor Impairs Parenchymal Plaque Development in an Alzheimer’s Disease Model”. Nature Communications 10 (1). 10.1038/s41467-019-11674-z.

Al-Onaizi, Mohammed A., Peter Thériault, Sarah Lecordier, Paul Prefontaine, Serge Rivest, and Ayman ElAli. 2022. “Early Monocyte Modulation by the Non-Erythropoietic Peptide ARA 290 Decelerates AD-like Pathology Progression”. Brain Behavior, and Immunity 99 (January): 363–82. 10.1016/j.bbi.2021.07.016.

Crapser, Joshua D, Joseph Ochaba, Neelakshi Soni, Jack C Reidling, Leslie M Thompson, and Kim N Green. 2019. “Microglial Depletion Prevents Extracellular Matrix Changes and Striatal Volume Reduction in a Model of Huntingtons Disease”. Brain 143 (1): 266–88. 10.1093/brain/awz363.

Sekiya, Hiroaki, Asato Tsuji, Yuki Hashimoto, Mariko Takata, Shunsuke Koga, Katsuya Nishida, Naonobu Futamura, et al. 2022. “Discrepancy between Distribution of Alpha-Synuclein Oligomers and Lewy-Related Pathology in Parkinson’s Disease”. Acta Neuropathologica Communications 10 (1). 10.1186/s40478-022-01440-6.

Wanner, IB, MA Anderson, B Song, J Levine, A Fernandez, Z Gray-Thompson, Y Ao, and MV Sofroniew. 2013. “Glial Scar Borders Are Formed by Newly Proliferated, Elongated Astrocytes That Interact to Corral Inflammatory and Fibrotic Cells via STAT3-Dependent Mechanisms after Spinal Cord Injury.”. J Neurosci 33: 12870–86.

Fernández-Klett, Francisco, and Josef Priller. 2014. “The Fibrotic Scar in Neurological Disorders”. Brain Pathology 24 (4): 404–13. 10.1111/bpa.12162.

Dzyubenko, Egor, Daniel Manrique-Castano, Christoph Kleinschnitz, Andreas Faissner, and Dirk M. Hermann. 2018. “Role of Immune Responses for Extracellular Matrix Remodeling in the Ischemic Brain”. Therapeutic Advances in Neurological Disorders 11 (January): 175628641881809. 10.1177/1756286418818092.

Dzyubenko, Egor, Daniel Manrique-Castano, Matthias Pillath-Eilers, Paraskevi Vasileiadou, Jacqueline Reinhard, Andreas Faissner, and Dirk M Hermann. 2022. “Tenascin-C Restricts Reactive Astrogliosis in the Ischemic Brain”. Matrix Biology 110 (June): 1–15. 10.1016/j.matbio.2022.04.003.

Dias, David O., Jannis Kalkitsas, Yildiz Kelahmetoglu, Cynthia P. Estrada, Jemal Tatarishvili, Daniel Holl, Linda Jansson, et al. 2021. “Pericyte-Derived Fibrotic Scarring Is Conserved across Diverse Central Nervous System Lesions”. Nature Communications 12 (1). 10.1038/s41467-021-25585-5.

Soderblom, Cynthia, Xueting Luo, Ezra Blumenthal, Eric Bray, Kirill Lyapichev, Jose Ramos, Vidhya Krishnan, et al. 2013. “Perivascular Fibroblasts Form the Fibrotic Scar after Contusive Spinal Cord Injury”. The Journal of Neuroscience 33 (34): 13882–87. 10.1523/jneurosci.2524-13.2013.

Göritz, Christian, David O. Dias, Nikolay Tomilin, Mariano Barbacid, Oleg Shupliakov, and Jonas Frisén. 2011. “A Pericyte Origin of Spinal Cord Scar Tissue”. Science 333 (6039): 238–42. 10.1126/science.1203165.

Bukkuri, Anuraag, Noemi Andor, and Isabel K. Darcy. 2021. “Applications of Topological Data Analysis in Oncology”. Frontiers in Artificial Intelligence 4 (April). 10.3389/frai.2021.659037.

Singh, Yashbir, Colleen M. Farrelly, Quincy A. Hathaway, Tim Leiner, Jaidip Jagtap, Gunnar E. Carlsson, and Bradley J. Erickson. 2023. “Topological Data Analysis in Medical Imaging: Current State of the Art”. Insights into Imaging 14 (1). 10.1186/s13244-023-01413-w.

Masoomy, Hosein, Behrouz Askari, Samin Tajik, Abbas K. Rizi, and G. Reza Jafari. 2021. “Topological Analysis of Interaction Patterns in Cancer-Specific Gene Regulatory Network: Persistent Homology Approach”. Scientific Reports 11 (1). 10.1038/s41598-021-94847-5.

Bhaskar, Dhananjay, William Y. Zhang, and Ian Y. Wong. 2021. “Topological Data Analysis of Collective and Individual Epithelial Cells Using Persistent Homology of Loops”. Soft Matter 17 (17): 4653–64. 10.1039/d1sm00072a.

Bonilla, Luis L., Ana Carpio, and Carolina Trenado. 2020. “Tracking Collective Cell Motion by Topological Data Analysis”. Edited by Philip K. Maini. PLOS Computational Biology 16 (12): e1008407. 10.1371/journal.pcbi.1008407.

Dawson, Madeleine, Carson Dudley, Sasamon Omoma, Hwai-Ray Tung, and Maria-Veronica Ciocanel. 2022. “Characterizing Emerging Features in Cell Dynamics Using Topological Data Analysis Methods”. Mathematical Biosciences and Engineering 20 (2): 3023–46. 10.3934/mbe.2023143.

Yamanashi, Takehiko, Mari Kajitani, Masaaki Iwata, Kaitlyn J. Crutchley, Pedro Marra, Johnny R. Malicoat, Jessica C. Williams, et al. 2021. “Topological Data Analysis (TDA) Enhances Bispectral EEG (BSEEG) Algorithm for Detection of Delirium”. Scientific Reports 11 (1). 10.1038/s41598-020-79391-y.

Saggar, Manish, James M. Shine, Raphaël Liégeois, Nico U. F. Dosenbach, and Damien Fair. 2022. “Precision Dynamical Mapping Using Topological Data Analysis Reveals a Hub-like Transition State at Rest”. Nature Communications 13 (1). 10.1038/s41467-022-32381-2.

Das, Soumya, D. Vijay Anand, and Moo K. Chung. 2023. “Topological Data Analysis of Human Brain Networks through Order Statistics”. Edited by Chad M. Topaz. PLOS ONE 18 (3): e0276419. 10.1371/journal.pone.0276419.

Al-Kofahi, Yousef, Alla Zaltsman, Robert Graves, Will Marshall, and Mirabela Rusu. 2018. “A Deep Learning-Based Algorithm for 2-D Cell Segmentation in Microscopy Images”. BMC Bioinformatics 19 (1). 10.1186/s12859-018-2375-z.

Greenwald, Noah F., Geneva Miller, Erick Moen, Alex Kong, Adam Kagel, Thomas Dougherty, Christine Camacho Fullaway, et al. 2021. “Whole-Cell Segmentation of Tissue Images with Human-Level Performance Using Large-Scale Data Annotation and Deep Learning”. Nature Biotechnology 40 (4): 555–65. 10.1038/s41587-021-01094-0.

Stringer, Carsen, Tim Wang, Michalis Michaelos, and Marius Pachitariu. 2020. “Cellpose: a Generalist Algorithm for Cellular Segmentation”. Nature Methods 18 (1): 100–106. 10.1038/s41592-020-01018-x.

Kruschke, JK, and TM Liddell. 2018. “Bayesian Data Analysis for Newcomers.”. Psychon Bull Rev 25: 155–77.

