## Supplementary material for "Dissecting glial scar formation by spatial point pattern and topological data analysis"

#### Supplementary Methods

Middle Cerebral Artery Occlusion (MCAO) surgery technique: 10.5281/zenodo.3559570

Intracardiac perfusion and brain fixation for immunohistochemistry: 10.17504/protocols.io.yxmvmk94og3p/v1

Staining of Gfap, Iba1, and NeuN on PFA-fixed mouse brain sections: 10.17504/protocols.io.4r3l27q5pg1y/v1

#### Supplementary Figures

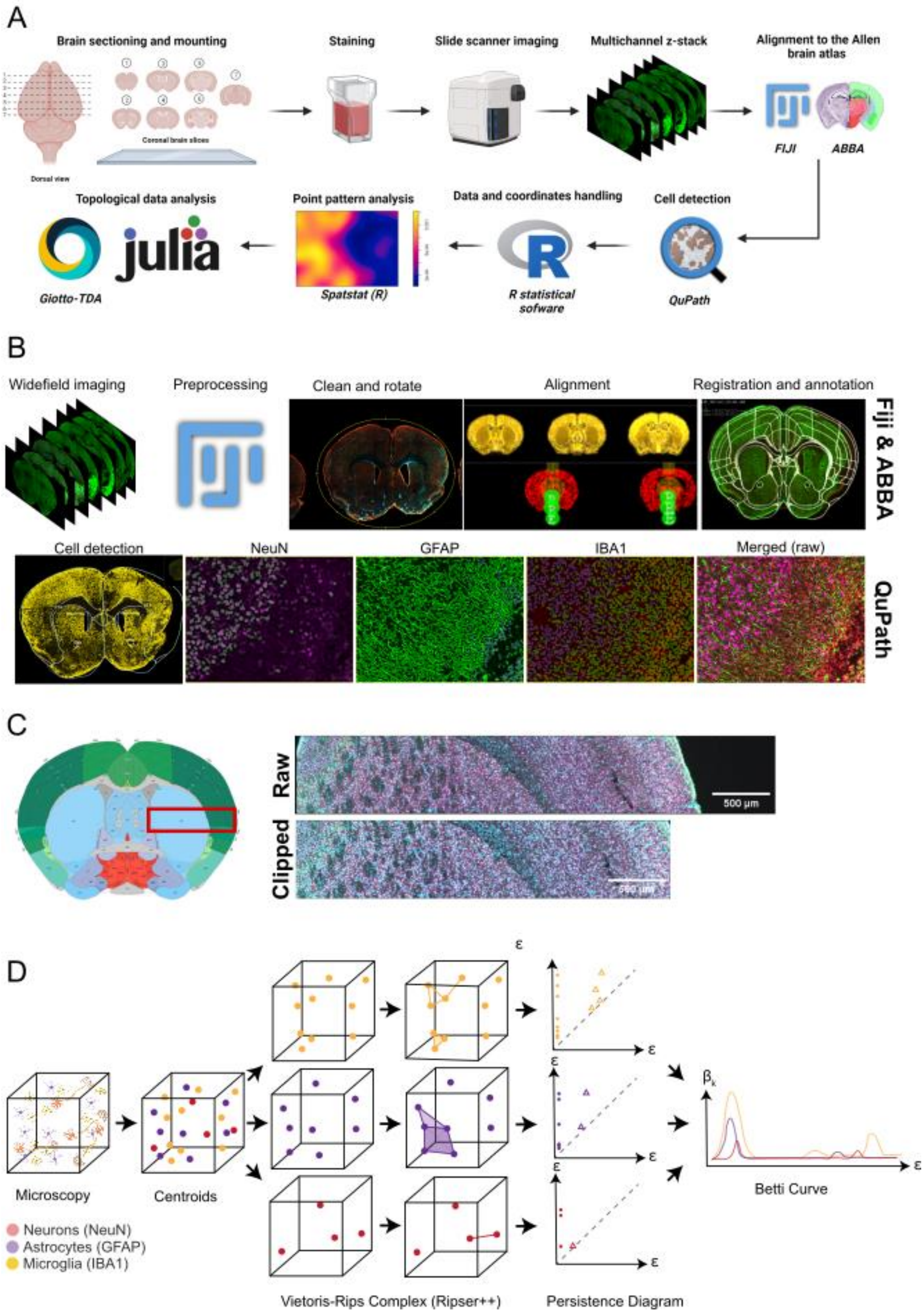

A
Brain sectioning and mounting
Coronal brain slices
Dorsal view
Staining
Slide scanner imaging
Multichannel z-stack
Alignment to the Allen brain atlas
FIJI
ABBA
Cell detection
QuPath
Data and coordinates handling
R statistical sofware
Point pattern analysis
Spatstat (R)
Topological data analysis
julia
Giotto-TDA
B
Widefield imaging
Preprocessing
Clean and rotate
Alignment
Registration and annotation
Fiji & ABBA
Cell detection
NeuN
GFAP
IBA1
Merged (raw)
QuPath
C
Raw
500 µm
Clipped
D
Microscopy
Centroids
Neurons (NeuN)
Astrocytes (GFAP)
Microglia (IBA1)
Vietoris-Rips Complex (Ripser++)
Persistence Diagram
Betti Curve

**Supplementary Figure 1. Workflow and analysis pipeline. A)** Workflow from brain sectioning to topological data analysis (schematic prepared with Biorender). **B)** Analysis pipeline using FIJI and ABBA (upper row) and QuPath (bottom row). Whole brain sections were imaged at 5x magnification and preprocessed to facilitate brain alignment and cell detection. After alignment and registration to the mouse Allen Brain Atlas, the brain sections are loaded in a QuPath project to perform unbiased quantification on NeuN-, GFAP-, and IBA1-expressing cells. **C)** Tiles at 10x magnification were acquired from the ventricular area to the border of the dorsolateral cortex (red rectangle). The images were manually clipped to adjust the observation window for PPA. **D)** (I) A three-dimensional image is transformed into a point pattern. (ii) Cell types are segregated and transformed into a simplicial complex by expanding a radius r ball around each point and connecting with edges when two balls intersect. (iii) Persistence diagrams for dimension 0 of data in (iV). Each dot represents a topological feature and its lifetime, i.e. persistence, in the filtration. (V). The Betti curve is a single representation of the topological features of each cell type.

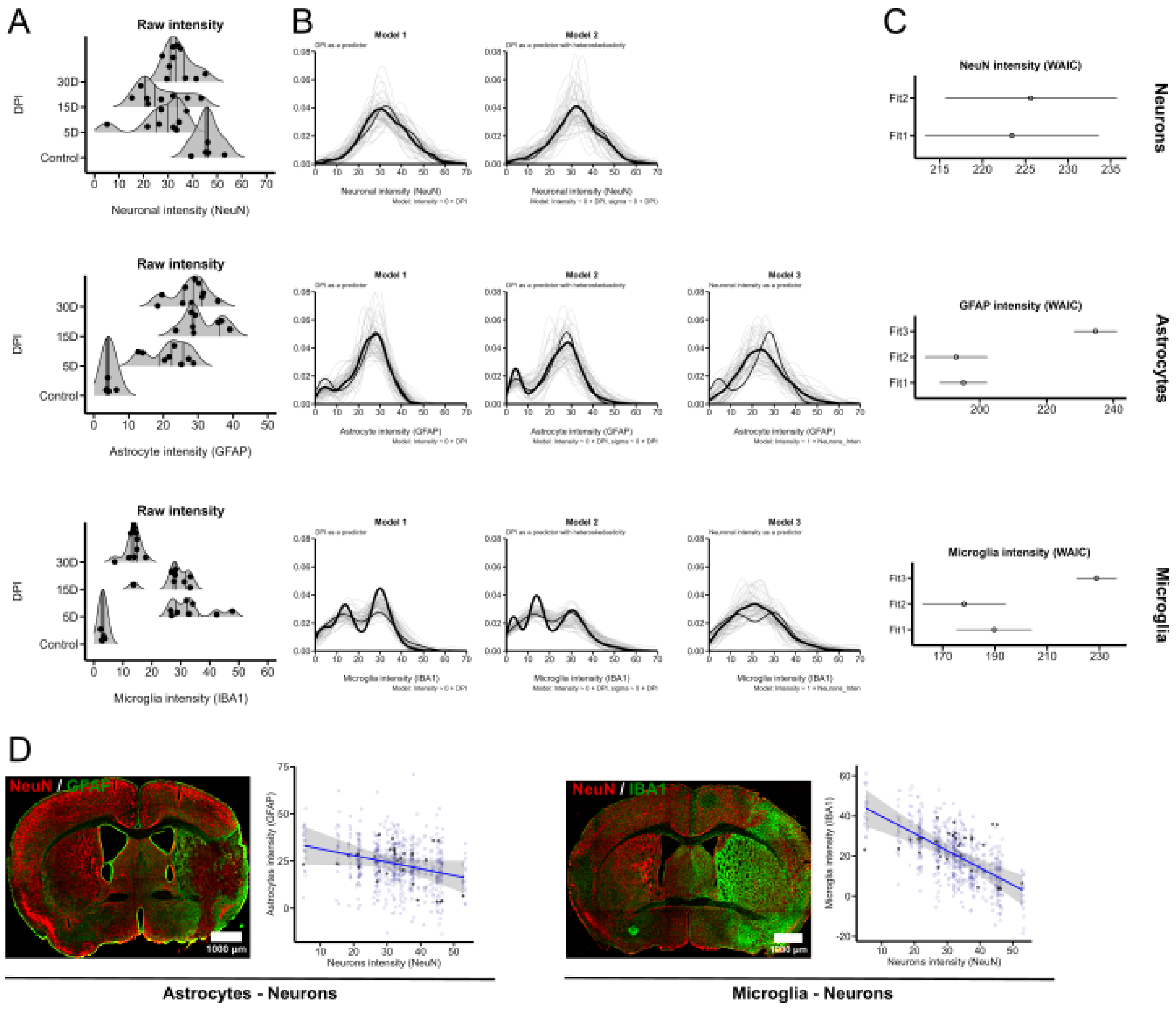


**Supplementary Figure 2. NeuN, GFAP and IBA1 intensity. A)** Density plots (ggridges) showing estimated spatial intensity values for neurons (NeuN), astrocytes (GFAP) and microglia (IBA1). Lines depict first, second, and third quartile, and black dots raw data points. **B)** Posterior predictive checks (brms) for fitted models. Faint lines show 50 random draws from the posterior distribution. The thin line is the posterior average, and the thick line is the observations. The model formulas are displayed as captions. **C)** Watanabe-Akaike information criterion (WAIC) of fitted models for NeuN, GFAP and IBA1. A higher value indicates a higher penalty to the model and lower out-of-sample prediction accuracy. **D)** Representative images of NeuN/GFAP and NeuN/IBA1 with the corresponding correlation between the markers' spatial intensity (all DPIs).

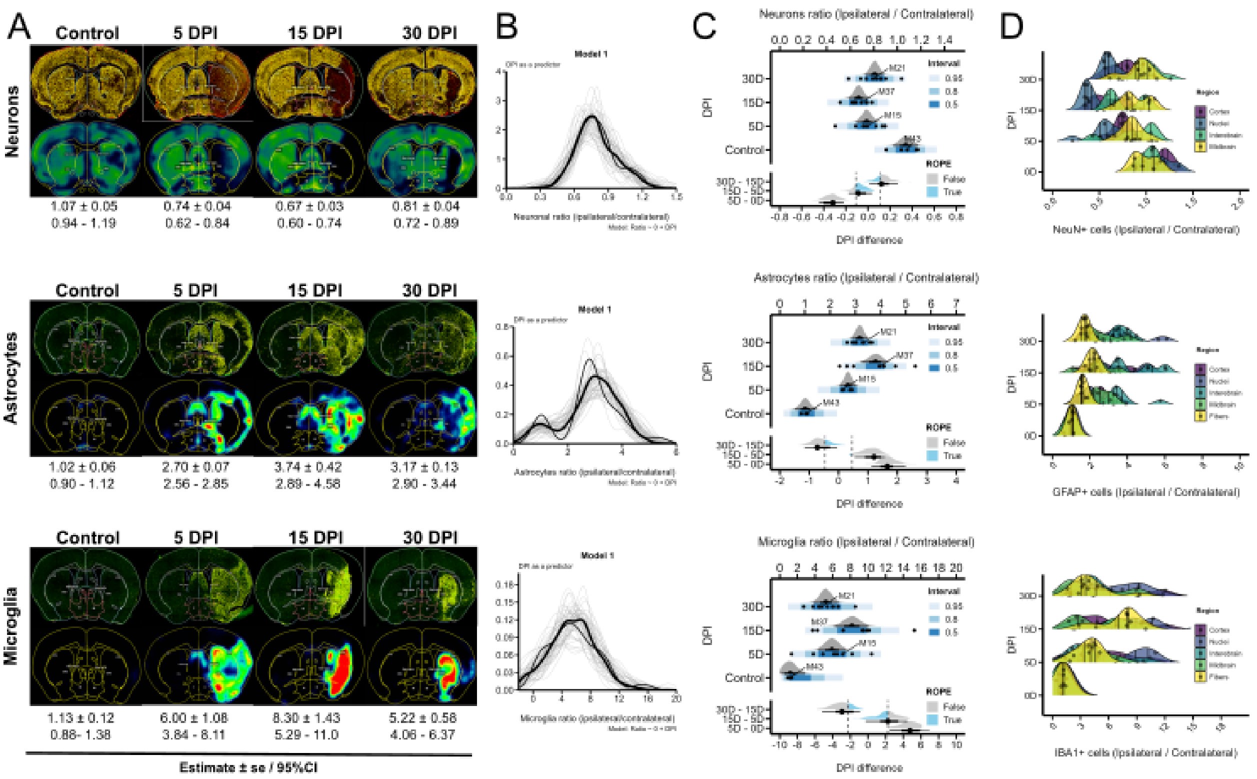


**Supplementary Figure 3. NeuN, GFAP and IBA1 cell ratios. A)** Sample images of Neurons (NeuN), astrocytes (GFAP), and microglia (IBA1). The upper row shows cell detections in QuPath (yellow) and the lower rows depict corresponding density maps. Estimates, standard deviation, and 95% confidence intervals for cell ratios (ipsilateral/contralateral) are shown per time point. **B)** Posterior predictive checks (brms) for each cell type corresponding to a model with DPI as a single predictor (see caption). Faint lines show 50 random draws from the posterior distribution. The thin line is the posterior average, and the thick line is the observations. **C)** Composite graphs showing Bayesian posterior distributions of cell ratios (spatial intensity) for neurons (NeuN), astrocytes (GFAP), and microglia (IBA1). Mean point estimates and their uncertainty are plotted as half-eye and point intervals. Raw data (black dots) is accompanied by prediction intervals (Brewer scale). The contrast between time points is plotted at the bottom of each composited graph. The region of practical equivalence (ROPE) encloses the values that fall within the time point variance. **D)** Density plots (ggridges) showing cell ratios per brain region. Lines depict first, second, and third quartile, and black dots raw data points.

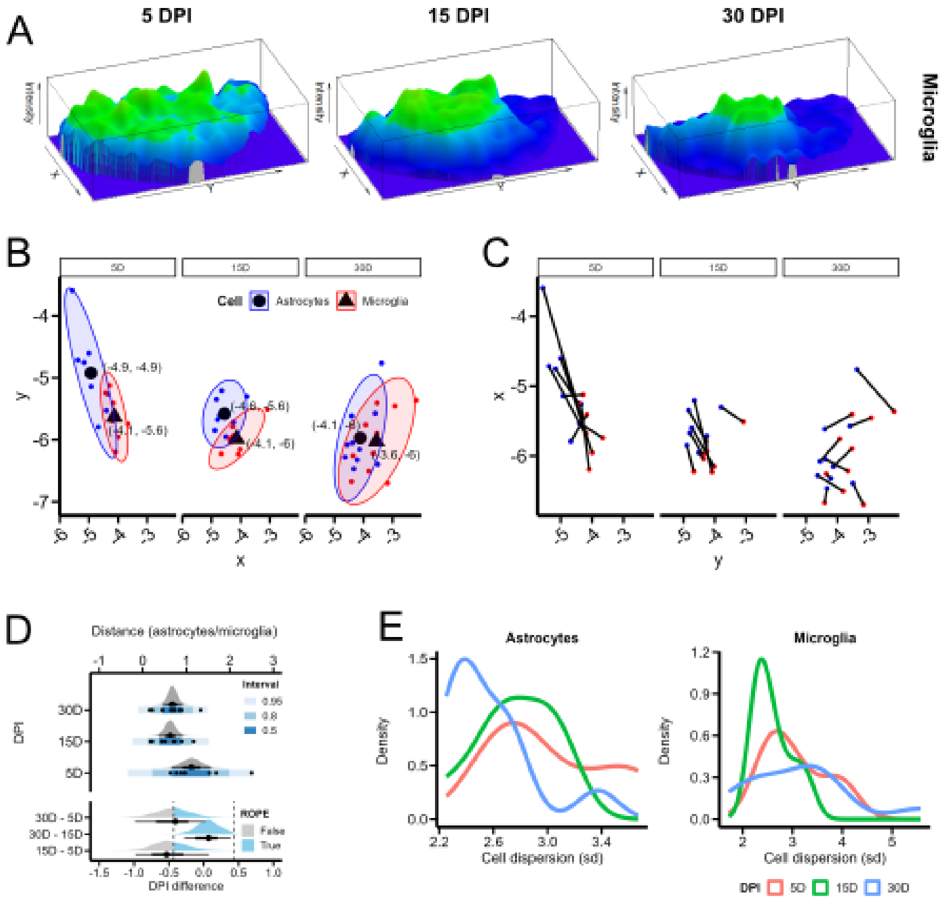


**Supplementary Figure 4. Distance/Dispersion measurements. A)** Perspective plots of superimposed brains showing microglia intensity (topo colors). The values decrease as the lesion progresses as part of the resolution of the inflammatory cascades. B) Scatter plot (ggplot) showing centroids for astrocytes (blue) and microglia (red) per DPI. Large marks depict the mean with corresponding xy coordinates and ellipses intervals at 0.80. **C)** Scatter plot (ggplot) showing connected lines between astrocytes (blue) and microglia (red) centroids per DPI. **D)** Posterior contrast between time points for astrocyte-microglia centroid distance. Mean point estimates and their uncertainty are plotted as half-eye and point intervals. The region of practical equivalence (ROPE) encloses the values that fall within the time point variance and are therefore considered equivalent to the null for robust estimation. **E)** Densities for astrocytes and microglia dispersion (sd) per DPI.

### Supplementary Tables

**Supplementary Table 1. Brain slice coordinates**. Bregma coordinates of the brain slices (30 µm coronal cuts) used in the present study. Section 1 (anterior) - Section 7 (posterior).

| Section | 1 | 2 | 3 | 4 | 5 | 6 | 7 |
|---|---|---|---|---|---|---|---|
| Bregma | 2.14 - 1.64 | 1.04 - 0.62 | 0.44 - - 0.06 | -0.18 - - 1.05 | -1.25 - - 1.85 | -1.95 - - 2.48 | -3.28 - 2.55 |
| Coronal level | 33 - 38 | 44 - 48 | 50 - 55 | 56 - 65 | 67 - 73 | 74 - 79 | 80 - 86 |

**Supplementary Table 2. Scanning parameters at 5x magnification.** Parameters used for scanning whole brain sections at 5x magnitude using an AxioScan Z1 slide scanner. Imaging was performed on NeuN, IBA1, GFAP, and DAPI-stained sections.

| | | | **Neun** | **IBA1** | **GFAP** | **DAPI** |
|---|---|---|---|---|---|---|
| **Objective** | Fluar 5x/0.25 M27 | **Channel** | AF647 | Cy3 | AF488 | DAPI |
| **Scaling per pixel** | 1.300 x 1.300 µm | **Excitation** | 653 | 458 | 493 | 353 |
| **Bit depth** | 16 bit | **Emission** | 668 | 561 | 517 | 465 |
| **File type** | Carl Zeiss Image (*.czi) | **Exposure** | 3 ms | 4 ms | 1 ms | 50 ms |

**Supplementary Table 3. Scanning parameters at 10x magnification.** Parameters used for scanning brain sections at 5x magnitude using an Zeiss Axio Observer.Z1 inverted epifluorescence microscope . Imaging was performed on NeuN, IBA1, GFAP, and DAPI-stained sections.

| | | | **Neun** | **IBA1** | **GFAP** | **DAPI** |
|---|---|---|---|---|---|---|
| **Objective** | ECM paln-NeoFluar 10x/0.30 M27 | **Channel** | AF647 | Cy3 | AF488 | DAPI |
| **Scaling per pixel** | 0.45 x 0.45 µm | **Excitation** | 653 | 458 | 493 | 353 |
| **Bit depth** | 16 bit | **Emission** | 668 | 561 | 517 | 465 |
| **File type** | Carl Zeiss Image (*.czi) | **Exposure** | 200 s | 250 s | 100 s | 10 ms |

**Supplementary Table 4. Estimation of spatial intensity**. Posterior estimates for the spatial intensity of NeuN, GFAP, and IBA1 in the whole ischemic hemisphere. The models (*Intensity ~ 0 + DPI)* were fitted using a student-t distribution. *Est.* (estimate), *se* (Standard error) *95% CI* (Credible intervals). See QN, Section 2 for posterior predictive checks and model diagnostics.

| **DPI** | **NeuN** | | | **GFAP** | | | **IBA1** | | |
|---|---|---|---|---|---|---|---|---|---|
| | **Est.** | **se** | **95% CI** | **Est.** | **se** | **95% CI** | **Est.** | **se** | **95% CI** |
| 0 | 45.5 | 3.2 | 38.7 – 52.3 | 4.6 | 2.1 | 0.9 – 9.25 | 3.2 | 1.2 | 0.6 – 6.5 |
| 5 | 28.4 | 2.9 | 22.4 – 34 | 21.9 | 1.8 | 18.1 – 25.6 | 31.9 | 1.8 | 28.3 – 36.1 |
| 15 | 36.7 | 2.9 | 20.9 – 32.7 | 31.1 | 1.8 | 27.4 – 34.8 | 28.8 | 1.4 | 25.7 – 31.8 |
| 30 | 34.2 | 2.3 | 29.5 – 39.1 | 27.7 | 1.6 | 24.3 – 30.9 | 13.8 | 1 | 11.4 – 16.1 |
| **Obs.** | 31 | | | 31 | | | 31 | | |
| **R2** | 0.42 | | | 0.76 | | | 0.81 | | |

**Supplementary Table 5. Cell ratio (ipsilateral/contralateral)**. Posterior estimates for the cell ratio (ipsilateral/contralateral) of NeuN, GFAP, and IBA1. The models (Ratio ~ *0 + DPI)* were fitted using a student-t distribution. *Est.* (estimate), *se* (Standard error) *95% CI* (Credible intervals). See QN, Section 3 for posterior predictive checks and model diagnostics.

| **DPI** | **NeuN** | | | **GFAP** | | | **IBA1** | | |
|---|---|---|---|---|---|---|---|---|---|
| | **Est.** | **se** | **95% CI** | **Est.** | **se** | **95% CI** | **Est.** | **se** | **95% CI** |
| 0 | 1.0 | 0.05 | 0.9 – 1.1 | 1.0 | 0.21 | 0.5 – 1.4 | 1.1 | 0.88 | -0.6 – 2.1 |
| 5 | 0.7 | 0.04 | 0.6 – 0.8 | 2.7 | 0.17 | 2.3 – 3.0 | 5.9 | 1.80 | 4.3 – 7.4 |
| 15 | 0.6 | 0.04 | 0.5 – 0.7 | 3.7 | 0.26 | 3.2 – 4.2 | 8.1 | 1.96 | 6.2 – 10.0 |
| 30 | 0.8 | 0.04 | 0.7 – 0.8 | 3.1 | 0.16 | 2.8 – 3.4 | 5.2 | 0.70 | 3.8 – 6.6 |
| **Obs.** | 31 | | | 31 | | | 31 | | |
| **R2** | 0.58 | | | 0.73 | | | 0.48 | | |

**Supplementary Table 6. Relative distribution of glia to neurons (mppm models)**. Spatial intensity of $GFAP^+$ and $IBA1^+$ cells conditional on NeuN (neuronal) spatial intensity. The *mppm* model (GFAP/IBA1 ~ NeuN_Dens, random = ~ NeuN_Dens DPI) shows estimates for fixed (population level) and random (group level) effects on the log scale. *Int* (Intercept), *Slope* (NeuN_Dens).

| **GFAP/NeuN** | | | **IBA1/NeuN** | | |
|---|---|---|---|---|---|
| **Fixed** | **Int(log)** | **Slope(log)** | **Fixed** | **Int(log)** | **Slope(log)** |
| | 3.18 | 0.002 | | 3.69 | -0.02 |
| **Random** | | | **Random** | | |
| 5 DPI | -0.51 | 0.01 | 5 DPI | 0.21 | -0.006 |
| 15 DPI | 0.19 | -0.002 | 15 DPI | 0.04 | 0.001 |
| 30 DPI | 0.32 | -0.008 | 30 DPI | -0.26 | -0.007 |

**Supplementary Table 7. Glial counts in tessellated neuronal regions.** Posterior estimates for $GFAP^+$ and $IBA1^+$ cell counts in tessellated (quantiles = 0, 20, 150) regions depicting low and high spatial intensity for neurons. The models ( *GFAP/IBA1 ~ 0 + NeuN_Dens : DPI )* were fitted using a student-t distribution. *Est.* (estimate), *se* (Standard error) *95% CI* (Credible intervals). See QN, Section 5 for posterior predictive checks and model diagnostics.

| | **GFAP/NeuN** | | | | **IBA1/NeuN** | | | |
|---|---|---|---|---|---|---|---|---|
| **DPI** | **NeuN_Dens** | **Est.** | **se** | **95% CI** | **NeuN_Dens** | **Est.** | **se** | **95% CI** |
| 5 | Low | 193 | 69 | 59 - 333 | Low | 484 | 68 | 349 - 623 |
| | High | 504 | 75 | 352 - 647 | High | 557 | 58 | 437 - 666 |
| 15 | Low | 410 | 66 | 277 - 537 | Low | 605 | 57 | 486 - 714 |
| | High | 503 | 81 | 344 - 662 | High | 238 | 56 | 129 - 353 |
| 30 | Low | 283 | 59 | 167 - 400 | Low | 225 | 42 | 140 - 310 |
| | High | 552 | 63 | 429 - 677 | High | 187 | 41 | 140 - 310 |
| **Sigma** | | 183 | 27 | 126 - 236 | | 126 | 24 | 84 - 181 |

**Supplementary Table 8. Relative distribution of GFAP$^+$ and IBA1$^+$ cells to the x-axis**. Spatial intensity of GFAP$^+$ and IBA1$^+$ cells conditional on hemispheric coordinates. The *mppm* model (GFAP/IBA1 ~ x, random = ~ x DPI ) shows estimates for fixed (population level) and random (group level) effects on the log scale. *Int* (Intercept), *Slope* (x-coordintes).

| | **GFAP** | | | **IBA1** | |
|---|---|---|---|---|---|
| **Fixed** | **Int(log)** | **Slope(log)** | **Fixed** | **Int(log)** | **Slope(log)** |
| | 3.2 | -0.007 | | 4.1 | 0.2 |
| **Random** | **Int(log)** | **Slope(log)** | **Random** | **Int(log)** | **Slope(log)** |
| 5 DPI | -1.1 | -0.2 | 5 DPI | -0.07 | -0.10 |
| 15 DPI | 0.3 | 0.02 | 15 DPI | -0.03 | -0.05 |
| 30 DPI | 0.8 | 0.01 | 30 DPI | 0.1 | 0.1 |

**Supplementary Table 9. GFAP/IBA1 covariance (mppm) in the ischemic hemisphere**. Spatial intensity of GFAP$^+$ cells conditional on the spatial intensity (density kernel) of IBA1$^+$ cells. The *mppm* model with interaction (GFAP ~ 0 + DPI * Miroglia_Dens) shows estimates for fixed (population level) on the log scale. *Est* (Estimate), se (standard error).

| **Coefficient** | **Est(log)** | **se(log)** |
|---|---|---|
| 5D | 3.1 | 0.03 |
| 15D | 3.1 | 0.02 |
| 30D | 2.9 | 0.02 |
| IBA1_Dens | -0.002 | 0.0009 |
| 15D:IBA1_Dens | 0.01 | 0.001 |
| 30D:IBA1_Dens | 0.02 | 0.001 |

**Supplementary Table 10. GFAP/IBA1 covariance (Tesselations) in the ischemic hemisphere.** Posterior estimates for GFAP⁺ cell counts in tessellated (quantiles = 0, 20, 150) regions depicting low and high spatial intensity for microglia. The model ( *GFAP ~ 0 + IBA1_Dens : DPI )* was fitted using a student-t distribution. *Est.* (estimate), *se* (Standard error) *95% CI* (Credible intervals). See QN, Section 6 for posterior predictive checks and model diagnostics.

| | **GFAP/IBA1** | | | |
|---|---|---|---|---|
| **DPI** | **IBA1_Dens** | **Est.** | **se** | **95% CI** |
| 5 | Low | 212 | 53 | 107 - 317 |
| | High | 482 | 57 | 371 - 597 |
| 15 | Low | 306 | 56 | 197 - 417 |
| | High | 616 | 55 | 506 - 727 |
| 30 | Low | 468 | 45 | 377 - 559 |
| | High | 366 | 46 | 276 - 458 |
| **Sigma** | | 141 | 18 | 106 - 179 |

**Supplementary Table 11. Relative distribution of astrocytes conditional on distance to microglia.** Spatial intensity of GFAP⁺ cells conditional on the distance (distfun) to IBA1⁺ cells. The *mppm* model with interaction (GFAP ~ DPI * Miroglia_Dist) shows estimates for fixed (population level) on the log scale.

| **Coefficient** | **Est.** | **se** | **CI** |
|---|---|---|---|
| Intercept | 7.4 | 0.01 | 7.43 - 7.48 |
| 15D | 0.8 | 0.02 | 0.79 - 0.86 |
| 30D | 1.32 | 0.02 | 1.29 - 1.36 |
| IBA1_Dist | 15.3 | 0.71 | 14.2 - 16.3 |
| 15D:IBA1_Dist | -2.7 | 1.18 | -4.48 - -1.08 |
| 30D:IBA1_Dist | -23.5 | 1.07 | -25.0 - -21.9 |

**Supplementary Table 12. Mean classification accuracy using 5-fold cross-validation to predict cell type from topological features using a feed-forward neural network.** Note that 100% accuracy in the recovery of cell type from the combination of dimension 0 (H0) and dimension 1 (H1) homology clearly indicates sufficient number of cells were sampled for persistent homology computation.

| **DPI** | $H_0$ | $H_1$ | $H_0 + H_1$ | **PCA(**$H_0$**)** | **PCA(**$H_1$**)** | **PCA(**$H_0 + H_1$**)** |
|---|---|---|---|---|---|---|
| [0.5ex] 5 | 1.00 | 1.00 | 1.00 | | | |
| 15 | 1.00 | 0.98 | 1.00 | | | |
| 30 | 1.00 | 0.97 | 1.00 | | | |

**Supplementary Table 13. Mean and standard deviation of classification accuracy using 3-fold cross-validation to predict DPI from topological features using a feed-forward neural network.** The accuracy of a random classifier is expected to be 0.25.

| **Cell Type** | **H0** | **H1** | **H0 + H1** |
|---|---|---|---|
| NeuN | 0.45 ± 0.02 | 0.36 ± 0.04 | 0.28 ± 0.11 |
| IBA1 | 0.24 ± 0.19 | 0.36 ± 0.05 | 0.36 ± 0.04 |
| GFAP | 0.26 ± 0.10 | 0.36 ± 0.04 | 0.27 ± 0.08 |
| NeuN+IBA1 | 0.51 ± 0.02 | 0.55 ± 0.02 | 0.52 ± 0.03 |
| IBA1+ GFAP | 0.38 ± 0.07 | 0.33 ± 0.03 | 0.48 ± 0.04 |
| NeuN+ GFAP | 0.29 ± 0.10 | 0.43 ± 0.09 | 0.33 ± 0.04 |
| NeuN+ IBA1+ GFAP | **0.71 ± 0.08** | 0.59 ± 0.02 | 0.77 ± 0.06 |

**Supplementary Table 14. Classification accuracy using unsupervised clustering to predict mouse identity from topological features.** The accuracy of a random classifier is expected to be 0.33.

| **Cell Type** | **H0** | **H1** | **H0 + H1** |
|---|---|---|---|
| NeuN | 0.79 | 0.38 | 0.70 |
| IBA1 | 0.82 | 0.41 | 0.79 |
| GFAP | 0.52 | 0.39 | 0.48 |
| NeuN+IBA1 | 0.79 | 0.36 | 0.68 |
| IBA1+GFAP | 0.86 | 0.44 | 0.71 |
| NeuN+GFAP | 0.88 | **0.49** | 0.71 |
| NeuN+IBA1+GFAP | **0.91** | 0.42 | **0.85** |
